# Crown defoliation decreases reproduction and wood growth in a marginal European beech population

**DOI:** 10.1101/474874

**Authors:** Sylvie Oddou-Muratorio, Cathleen Petit-Cailleux, Valentin Journé, Matthieu Lingrand, Jean-André Magdalou, Christophe Hurson, Joseph Garrigue, Hendrik Davi, Elodie Magnanou

## Abstract

1. Abiotic and biotic stresses related to climate change have been associated to increased crown defoliation, decreased growth and a higher risk of mortality in many forest tree species, but the impact of stresses on tree reproduction and forest regeneration remains understudied. At dry, warm margin of species distributions, flowering, pollination and seed maturation processes are expected to be affected by drought, late frost and other stresses, eventually resulting in reproduction failure. Moreover, inter-individual variations in reproductive performances versus other performances (growth, survival) could have important consequences on population’s dynamics.
2. We investigated the relationships between individual crown defoliation, growth and reproduction in a drought-prone population of European beech, *Fagus sylvatica*. We used a spatially explicit mating model and marker-based parentage analyses to estimate effective female and male fecundities of 432 reproductive trees, which were also monitored for basal area increment and crown defoliation over nine years.
3. Female and male fecundities markedly varied among individuals, more than did growth. Both female fecundity and growth decreased with increasing crown defoliation and competition and increased with size. Male fecundity only responded to competition, and decreased with increasing competition. Moreover, the negative effect of defoliation on female fecundity was size-dependent, with a slower decline in female fecundity with increasing defoliation for the large individuals. Finally, a trade-off between growth and female fecundity was observed in response to defoliation: some large trees maintained significant female fecundity at the expense of reduced growth in response to defoliation, while some other defoliated trees rather maintained high growth at the expense of reduced female fecundity.
4. *Synthesis*. Our results suggest that while decreasing their growth, some large defoliated trees still contribute to reproduction through seed production and pollination. This non-coordinated decline of growth and fecundity at individual-level in response to stress may compromise the evolution of stress-resistance traits at population level, and increase forest tree vulnerability.

## Introduction

The increasing impact of stresses associated with climate and global change are likely to cause widespread forest decline, eventually leading to massive tree mortality due to inability to recover from stresses (Allen et al., 2010; McDowell et al., 2011). Depending on their frequency, duration, or intensity, abiotic stresses (e.g. drought, wind throw, flood, heavy snow, late frosts, fire) and biotic stresses (predation, competition) have the potential to alter tree structure (e.g. branch breakage, leaf fall), physiological processes (e.g. hydraulic failure, reduced photosynthesis) and overall vigour (e.g., crown defoliation) and performances (e.g. reduced growth, reproduction and survival). Moreover, as stresses often co-occur and interact, it is notoriously difficult to disentangle the drivers of tree decline observed in a given environment. Hence, the diversity of both stresses and decline components needs to be accounted for in order to better predict forest decline in response to environmental change.

The warm and dry margins of tree species distributions are expected and already observed to suffer massive forest decline, driven by increasing temperatures, drought, late frost and other stresses. Most importantly, prolonged droughts and high temperatures have been extensively associated to decreasing tree growth and forest productivity (Zhao & Running, 2010; Zimmermann, Hauck, Dulamsuren, & Leuschner, 2015), increasing crown defoliation and leaf fall (Dobbertin, 2005; Galiano, Martínez-Vilalta, & Lloret, 2011) and higher risks of tree mortality (Adams et al., 2017; Allen et al., 2010; Anderegg, Kane, & Anderegg, 2013). There is also an increasing concern that the advance in spring phenology currently observed in many species expose them to a higher risk of late frost, with damaging effects on crown development (Bigler & Bugmann, 2018; Charrier, Ngao, Saudreau, & Améglio, 2015). Overall, the response of tree sexual reproduction and forest regeneration to abiotic and biotic stresses remain largely under-documented, despite the critical importance of reproduction for the maintenance, demography and adaptation of populations at the rear-edge of species distribution (Hampe & Petit, 2005). Here, we consider sexual reproduction globally, including all the stages from floral initiation to the production of mature seeds.

Based on species physiology, abiotic stresses such as droughts or late frosts are expected to directly reduce plant sexual reproduction through altered reproductive phenology (i.e. the timing of flowering and fruiting), a higher risk of pollen abortion or pollination failure, a shorter seed maturation cycle and/or a higher risk of seed abortion (Bykova, Chuine, Morin, & Higgins, 2012; Hedhly, Hormaza, & Herrero, 2009; Zinn, Tunc-Ozdemir, & Harper, 2010). Moreover, indirect negative effects are also expected: for instance, by decreasing photosynthetic activity, leaf fall may reduce the amount of stored resources to invest in reproduction of the next year (Obeso, 1988). The few results available so far from experiments manipulating stresses *in situ* overall support these expectation of negative impacts of stresses on reproduction and growth. By manipulating temperature during pollen dispersal and germination, Flores-Rentería *et al.* (2018) demonstrated negative impacts of high temperatures on male reproduction, particularly on pollen viability of *Pinus edulis*. Bykova *et al.* (2018) also showed that water deficit increases pollen abortion and thus decreases pollen production in *Quercus ilex*. In *Quercus ilex*, Pérez-Ramos *et al.* (2010) showed that reduced water availability increased the rate of acorn abortion, while Sanchez-Humanes & Espelta (2011) showed that increased drought reduces acorn production in coppice. In parallel, these drought-manipulation experiments demonstrated that growth generally decreases (e.g. Delaporte, Bazot, & Damesin, 2016; Lempereur et al., 2015) and crown defoliation increases (e.g. Galiano et al., 2011) in response to increasing water stress.

On the other hand, stresses have been hypothesised to shift patterns of resource allocation and act like a cue stimulating higher reproductive effort and reduced growth (Bréda, Huc, Granier, & Dreyer, 2006; Lee, 1988; Pulido et al., 2014; Wiley, Casper, & Helliker, 2017). However, results from experiments manipulating stresses *in situ* hardy support this expectation, even though they highlight flexible allocation rules to reproduction and growth in response to stress. Using experimental warming, Sherry *et al.* (2007) demonstrated divergent responses of reproductive phenology to water stress for grass species: some species advanced their flowering and fruiting phenology before the peak of summer heat while other species started flowering after the peak temperature and delayed their reproduction. In *Quercus ilex*, Pulido et al. (2014) did not find evidence that drought enhances resource allocation to reproduction but suggested that the negative individual correlation between growth and female reproduction observed in controlled conditions disappears under limited resources (including water stress). The clearest experimental evidence of the positive impact of biotic and abiotic stresses on reproduction can be found in the literature on (fruit) tree orchards. Among the cultural practices allowing early and abundant flowering, water stress is used to enhance flower initiation in conifers, while hot, dry summers are reported to induce abundant seed crops in both conifers and broadleaved species (Meilan, 1997). Another practice relies on circumferential girdles (the removal of a swath of the bark, down to the phloem, around the entire stem), which are associated to reduced vegetative growth and increased fruiting (Bonnet-Masimbert & Webber, 2012). Finally, pruning (the reduction of crown leaf area) is also recommended to favour reproductive development while reducing vegetative growth in fruit trees (Karimi et al., 2017).

Taken together, these results suggest that stress impacts on reproduction and the relationship between reproduction and growth in response to stress both need further investigations (Figure 1). Environment-manipulation experiments typically use a limited number of individuals in controlled conditions to characterize the fine impacts of stresses on the physiological mechanisms driving plant performances (eventually testing for individual effects, e.g. Camarero, Gazol, Sangüesa-Barreda, Oliva, & Vicente-Serrano, 2015). Besides these ecophysiological approaches, we also need population ecology approaches to investigate the among-individual variations in reproductive and vegetative performances in response to stress, and their consequences on population dynamics. Here, we propose to use crown defoliation taken as an indicator of stress, and to analyse the relationship between crown defoliation, reproduction and growth in order to test two hypotheses. First (H1), crown defoliation is associated to a proportional decrease in growth and reproduction through the impact of stresses on the resources allocated to these performances, so that the relationship between reproduction and growth does not change with increasing crown defoliation. Alternatively (H2), if defoliation or stresses act like a cue stimulating reproductive performances at the expense of reduced growth, then the relationship between reproduction and growth should change with increasing crown defoliation.

**Figure 1:**
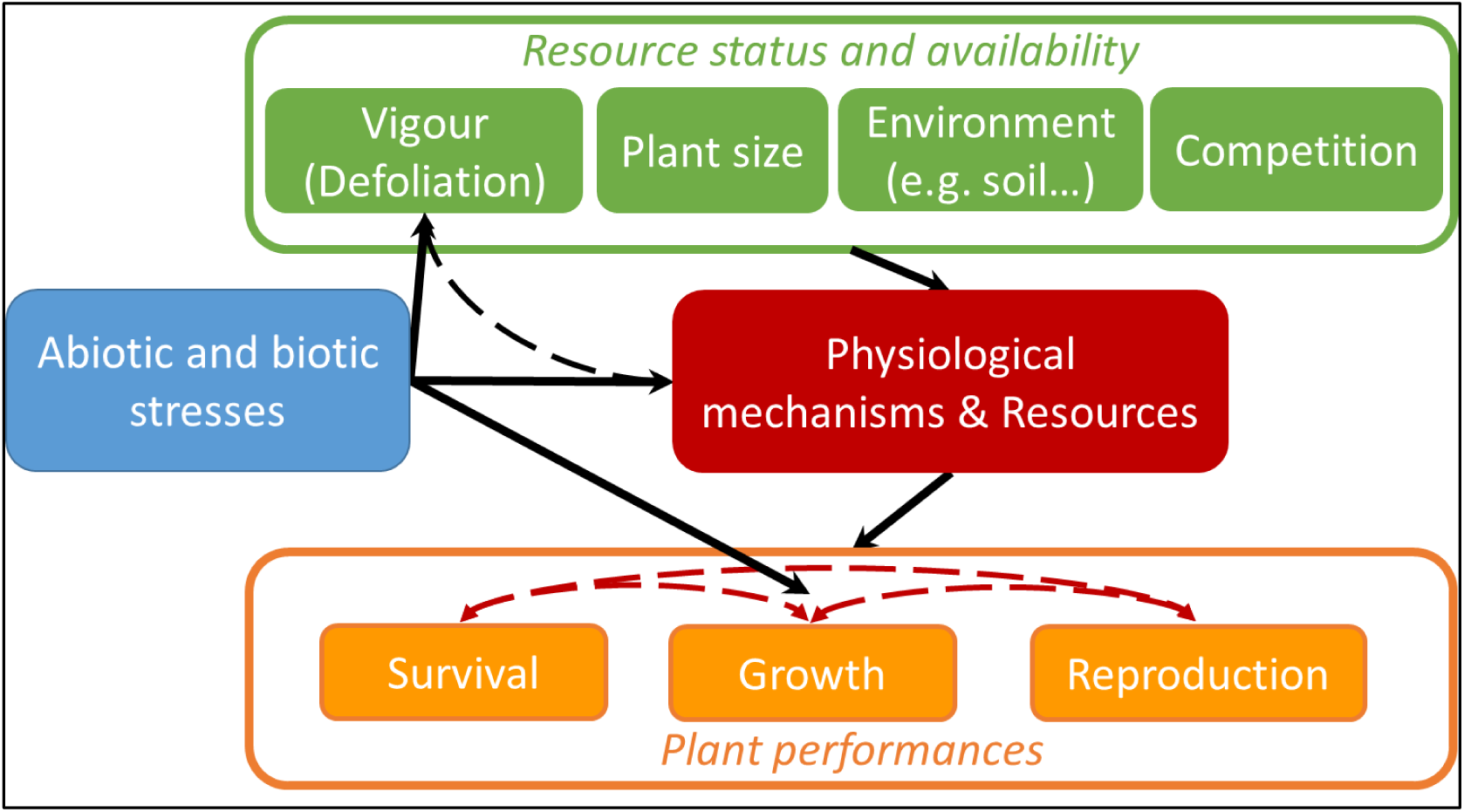
Conceptual framework for the impacts of abiotic and biotic stresses on individual plant performances (survival, growth, and reproduction). Stresses are expected to affect performances through their effect on the physiological mechanisms and the level of resources of the plant. Usually not easily measurable in natural populations, this level of resources also varies among individuals as a function of (i) plant resource status, a combination of plant size and vigour, and (ii) resource availability, which depend on the quality of the local environment and on competition. Vigour (eg crown defoliation) can in turn rapidly change in response to stress or to the level of resources (i.e. potential feedback loops, dashed arrows), and therefore it is itself an indicator of stress. Finally, stresses can act like cues changing resources allocation to survival, growth and reproduction, thereby affecting their correlations at individual level (eg., tradeoffs, dashed red arrows).

We focus here on the European beech (*Fagus sylvatica* L.), a major broadleaf tree species considered to be vulnerable to summer drought. Beech is a monoecious, wind-dispersed species, and shows an intermittent production of large seed crops synchronized across a population (i.e., masting), triggered both by weather and plant resource status (Vacchiano et al., 2017). Hacket-Pain, Lageard, & Thomas (2017) showed that drought years were associated to both reduced reproduction and growth, while during non-drought years, both masting and high growth could be observed. By contrast, Bréda *et al.* (2006) reported increased seed production associated to leaf fall in high drought years, even though this relationship between crown defoliation and fruit production may not be directly causal. This study investigates the relationships between tree defoliation, growth and fecundity in 432 individuals within a single, rear-edge natural population of *F. sylvatica* in Southern France, where crown defoliation and mortality are being surveyed since 2003 (Petit-Cailleux et al., submitted). We used molecular markers and parentage analyses to estimate effective, relative female and male fecundities, which integrate the success of pollination and germination processes cumulated from 2002 to 2012. Growth over the same period was assessed through inventory data, completed by ring-width measurements. We analysed the relationships between crown defoliation, growth and fecundity at the among-individual scale in order to (i) characterize the decline in fecundity and growth associated with defoliation and (ii) investigate the correlation between growth and fecundity in response to defoliation.

Our study is based on the well-accepted hypothesis that recurrent defoliation is related to physiological stresses and symptomatic of a declining health in beech (Bréda et al., 2006; Peñuelas & Boada, 2003). Using statistical models and 4327 trees individually surveyed, a companion study in the same population showed that crown defoliation increases the risk of mortality (Petit-Cailleux et al., submitted). Moreover, simulations with a process-based physiological model indicated that the mortality rate in this population is driven by a combination of drought-related processes (conductance loss, carbon reserve depletion) and late frost damages (Petit-Cailleux et al., submitted). Crown defoliation thus appears as an appropriate indicator of a higher intrinsic sensitivity to these stresses, and/or a higher impact of these stresses due to a lower availability of resources in a heterogeneous environment.

## Material and methods

### Study site

Located in southern France and bordering Spanish Cataluña, the Massane forest is situated on the foothills of Eastern Pyrenees (Figure 2B). This coastal range, called Albera massif, covers about 19,000 ha on the French territory. The Massane forest National Nature Reserve (42° 28’ 41” N, 3° 1’ 26” E) was created in 1973. It covers 336 ha on the highest part of the Massane valley, from 600 to 1,127 m a.s.l., and is only around 5 km far from the Mediterranean Sea. The site is under a meso Mediterranean climate influence (Quézel & Médail, 2003: mean annual temperature = 11.95°C; annual precipitations = 1164.9 mm, monitored on site since respectively 1976 and 1960; Figure S1A). This site is one of the French beech location the most prone to water stress (Figure S1B).

**Figure 2:**
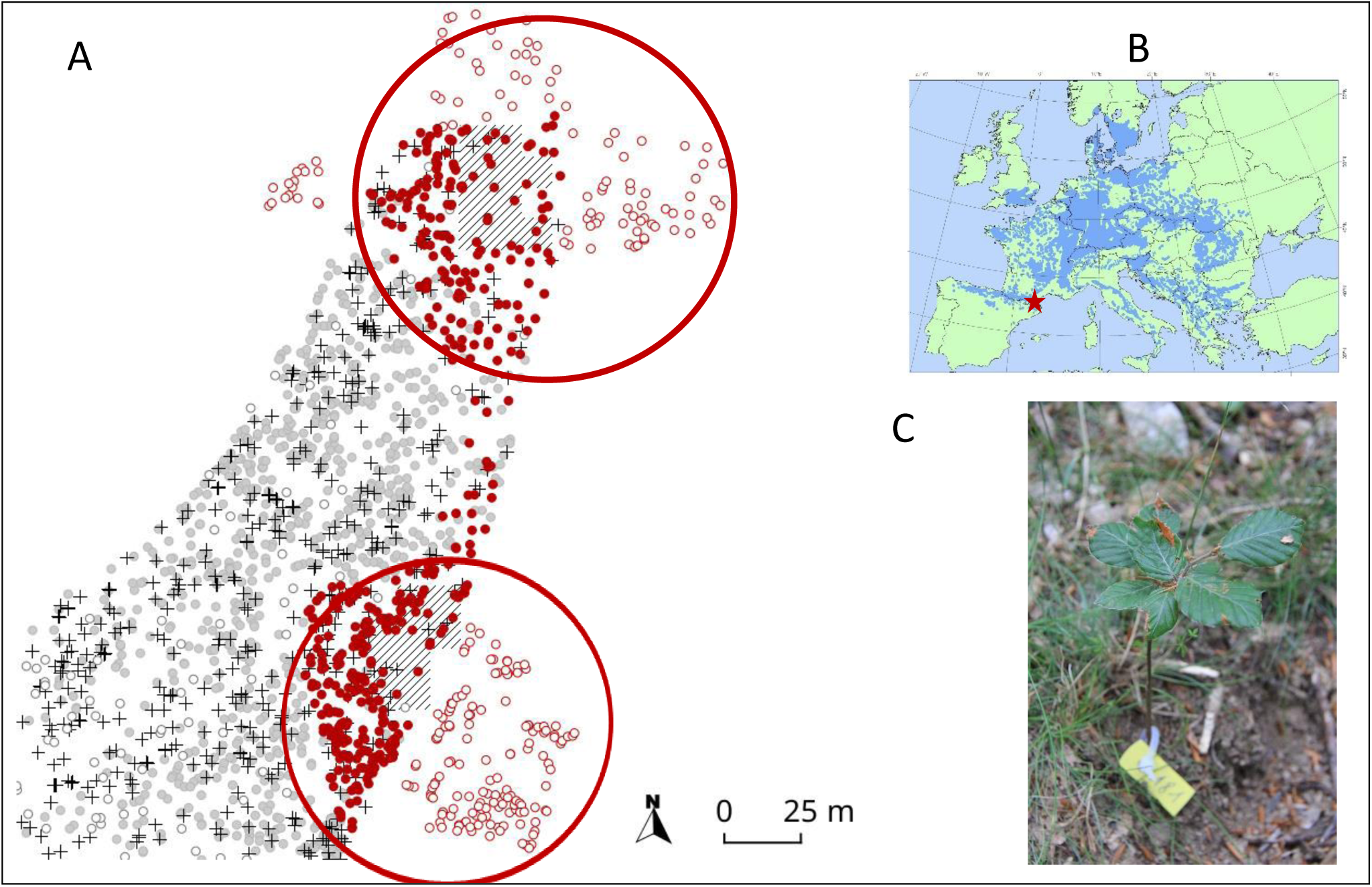
Study site and (A) sampling design: red filled dots (•) represent the 432 beech trees for which individual fecundity, growth and defoliation were assessed. Hatched squares represent the seedlings patches used to estimate fecundity through parentage analyses and mixed-effect mating models (MEMM). The two red circles encompass all the individuals (seedlings and adult) used for MEMM analyses. Red empty dots (○) represent the 244 beech trees outside of the protected area and included in the fecundity analyses (but not phenotyped for growth and defoliation). Grey dots (•) and crosses (+) represent other beeches within the protected area not included in the fecundity analyses either because they were far from sampled seedlings (•) or because they were dead in 2012 (+). Empty dots (○) represent other species within the protected area. (B) Study site location (red star) on European Beech distribution map (source Euforgen). (C) Picture of a sampled seedlings.

More than half of the Reserve is constituted of an old grown forest, where no logging operation has been performed since at least 1886. The canopy is dominated by European beech in mixture with downy oak (*Quercus pubescens* Willd.), maples (*Acer opalus* Mill., *Acer campestris* L., *Acer monspessulanum* L.) and holly (*Ilex aquifolium* L.). A 10 ha fenced plot was remote from any cow grazing since 1956. All trees from this protected plot are monitored since 2002.

### Adult seed-tree inventory and phenotyping

This study was conducted on two circular-shaped plots (as classically used in parentage analyses) covering 0.17 ha in total, where all the 683 alive adult beeches were mapped and collected for genetic analyses in 2012 (red dots on Figure 2). Although beech reproduction is mostly sexual, vegetative reproduction may occasionally occur, with the production of stump shoots resulting in multiple stems (i.e. several ramets for a single genet). In obvious cases of vegetative reproduction (ie root-connected stems), we sampled only the biggest ramet of each genet for genetic analyses.

Only 439 among the 683 collected beeches were included within the protected plot and monitored since 2002 (filled dots on Figure 2). The monitoring consisted first in measuring tree size as the diameter at breast height (DBH) in 2002 and 2012, which allowed us to derive the basal area (BA=π*DBH^2^/4). Individual growth was measured by the basal area increment (herein BAI) between 2002 and 2012, as estimated by: BAI=π(DBH^2^_2012_-DBH^2^_2002_)/4.

The presence of dead branches and leaves was recorded each year between 2004 and 2012 as a qualitative measure (1=presence; 0=absence). We used the sum of these nine annual defoliation scores (herein DEF) as an integrative, qualitative ordered measure, combining the recurrence of defoliation and the ability to recover from defoliation.

The conspecific local density (herein *Dens*_*dmax*_) was estimated as the number of beech neighbors found within a radius of *dmax* around each mother-tree. We also used the Martin-Ek index (Martin & Ek, 1984) to quantify the intensity of competition on a focal individual *i*. This index (herein *Compet*_*dmax*_) accounts simultaneously for the diameter and the distance of each beech competitor *j* to the competed individual *i*:

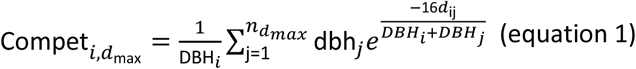

wher*e* DBH_*i*_ and DBH_*j*_ are the diameter at breast height (in cm) of the competed individua*l i* and of competito*r j* (any adult tree of any species with DBH_j_>DBH_i_), *n*_*dmax*_ the total number of competitors in a given radius *d*_*ma*x_ (in m) around each individual *i*, and *d*_*ij*_ the distance between individuals i and j. We computed a total of 20 *Dens*_*dmax*_ *0020*variables and 20 C*ompet* _*dmax*_ variables, by considering *d*_*max*_ values between 1 and 20 m with a 1m-step. The *Dens*_*dmax*_ variables were strongly and positively correlated with each other’s, and so were the C*ompet*_*dmax*_ variables, but *Dens*_*dmax*_ variables were not correlated with C*ompet* _*dmax*_ variables (Figure S2).

### Offspring sampling and genotyping

To estimate adult fecundity, we sampled 365 seedlings established amidst the 683 genotyped adult beeches (shaded quadrats on Figure 2). Two cohorts of seedlings were sampled exhaustively within a selected number of quadrats at the center of each circular plot: 165 “young” seedlings germinated in spring 2012 (masting year 2011), and 200 “old seedlings” germinated from spring 2011 back to spring 2001 (age was estimated using annual bud scars). Qualitative surveys indicated that masting occurred in years 2002, 2004, 2006 and 2009. In this study, the two seedlings cohorts were mixed, in order to estimate cumulated reproduction from 2001 to 2012.

The genotypes of the 683 alive adult beeches and 365 seedlings were scored at a combination of 18 microsatellite loci (Table S1). DNA extraction, PCR amplifications and genotype scoring with a MegaBACE 1000 sequencer were performed using the conditions described by Oddou-Muratorio, *et al.* (2018). The total number of alleles observed in each cohort was greater than 95 (Table S1). Adult genotypes revealed seven pairs of clones among the adult beeches. We checked that these clones were always spatially clustered, and kept only one ramet for each genet in the MEMM analyses (i.e. 676 adult beeches).

### Inference of male and female relative fecundities: MEMM analyses

Male and female fecundities were jointly estimated with the pollen and seed dispersal kernels in a Bayesian framework implemented in the MEMM program (Oddou-Muratorio et al., 2018). MEMM is one of the recently developed spatially explicit mating models based on naturally established seedlings (see also Burczyk, Adams, Birkes, & Chybicki, 2006; Goto, Shimatani, Yoshimaru, & Takahashi, 2006; Moran & Clark, 2011; Oddou-Muratorio & Klein, 2008). These models provide joint estimates of individual male and female fecundities together with the pollen and seed dispersal kernels and mating system parameters, so that estimates of fecundity are not biased by the confounding effects of spatial and sampling designs (the arrangement of male/female parents and sampled seedlings).

Briefly, the Mixed-Effect Mating Model (MEMM) considers that each sampled seedling originates either (i) from a mother tree located outside the study site (implying seed immigration) or (ii) from a mother tree located within the study site. The latter case includes three possible origins of the fertilizing pollen: (i) pollen immigration, (ii) selfing, or (iii) pollination by a male tree located within the study site. The approach bypasses parentage assignation and focuses instead on the fractional contribution of all adults, either as female or as male parent, to each seedling (see Appendix A1 for details). For instance, the probability *π*_*Sij*_ of each sampled female tree *j* to contribute to the seedling pool at the spatial location of seedling *i* is modeled as:

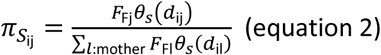

where *F*_*Fj*_ and *F*_*Fl*_ are the female fecundities of mother *j* and *l*, respectively; *d*_*ij*_ and *d*_*il*_ are the distances between seedling *i* and mother *j* and *l*, respectively; and *ϑ*_*s*_ is the seed dispersal kernel. Both the seed and pollen dispersal kernels (*ϑ*_*s*_ and *ϑ*_*p*_) are modelled using a power-exponential function. All the parameters of the model are estimated in a Bayesian framework (Appendix A1). Note that *F*_*F*_ (and *F*_*M*_) estimates are relative, with the average *F*_*F*_*-*value (and *F*_*M*_*-*) over the entire parent population fixed to 1.

Note that the fecundity estimates provided by MEMMs are related but not equivalent to the traditional resource-based estimates of female (i.e. the biomass/number of ovules, seeds, ovuliferous flowers or fruits) and male fecundity (i.e. the biomass/number of pollen grains or staminate flowers). First the latter estimate the resources allocated by each plant to reproduction while the former can only estimate a relative amount of pollen or seeds produced by each plant as compared to other plants. Second, the resource-based estimate is a pre-dispersal evaluation of seed and pollen production while MEMM estimates an effective amount of pollen achieving successful fertilization, and of seeds achieving successful germination. In consequence, MEMM-based estimates of fecundities account for individual effects (either maternal or genetic) that act independently on location to modify the success of mating, seed maturation or germination, or early survival during the post-dispersal processes preceding the sampling stage (Oddou-Muratorio et al., 2018).

For the estimation, we accounted for typing errors at microsatellite loci, with two possible types of mistyping: in the first type, the allele read differs only by one motif repeat from the true allele with a probability P_err1_, while in the second type, the allele read can be any allele observed at this locus with a probability P_err2_. We considered a mixture of the two error types, with P_err1_ = 0.01 and P_err2_ = 0.01. We ran 10 Markov chain Monte Carlo (MCMC) of 10,000 steps, each with additional 500 first MCMC steps as burn-in, checked that the different chains converged to the same value visually, and then combined the 10 chains together. Individual female (F♀) and male (F♂) fecundities were summarized by their median value across the 100,000 iterations.

### Adult subsampling for dendrochronological analyses

We selected 90 trees within the protected plot for which we sampled cores to measure ring-width. These 90 trees were chosen to represent contrast in terms of defoliation and female fecundity (Figure S3). Cores were extracted in February 2016 at 1.30 m above ground. After sanding, cores were scanned at high resolution (1200 dpi). Boundary rings were read using CooRecorder v 9.0. Ring width were transcribed, individual series were checked for missing rings and dating errors and mean chronologies were calculated using Cdendro 9.0 (CDendro 9.0 & CooRecorder 9.0; Cybis Elektronik & Data AB. Sweden). Using the sum of ring width increments between 2002 and 2012 (Σrw), the growth of the 90 individuals between 2002 and 2012 was estimated as: BAI_wood_=π((DBH_2002_/2+Σrw)^2^-DBH^2^_2002_/4).

### Statistical analyses of the ecological drivers of growth and fecundities

Our objective herein was to test whether defoliation significantly affected individual growth and female/male fecundity. For each response variable independently (i.e., growth as measured by BAI, and fecundities as estimated with MEMM), we considered the following initial linear model:

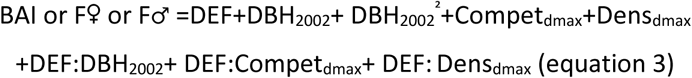

where all the predictors are quantitative variables (Table S2). Besides the target defoliation factor (DEF), this model includes one size-related factor (DBH_2002_), and two competition-related factors (Compet_dmax_ and Dens_dmax_). Size and competition are considered here as “nuisance” parameters, susceptible to blur the signal between defoliation, growth and reproduction. A quadratic effect of DBH_2002_ was also included, as growth and sometimes fecundity are known to be proportional to basal area. Density and competition index can both be relevant to capture competition effect on growth or fecundity, and moreover, their influence may vary with the distance up to which competitors are accounted for. Therefore, we first selected the best Compet_dmax_ and the best Dens_dmax_ terms for each response variable independently using the model described by equation 3 without interaction terms, and retaining the d_max_ values leading to the highest R^2^. Then, we included interactions terms (the three last terms in equation 3) to investigate specific effects of defoliation depending on individual size or on the level of competition.

The model was fitted on 432 focal adult beech trees within the protected plot (Figure 2) for which BAI was estimated from inventory data. All response variables were log-transformed to approach Gaussian distribution and to account for the higher variance associated to higher fecundity or higher growth. We visually inspected the relationship between each predictor and each response variable (Figure S4). For each response variable, we selected the most parsimonious model based on the AIC using the functions ‘lm’ and ‘step’ in R 3.3 (R Core Team 2018). The residuals were visually inspected through a plot of residuals vs predicted. Interaction effects were visualized with the package ‘jtools’ (Long, 2018).

Collinearity resulting from correlations between predictor variables is expected to affect statistical significance of correlated variables by increasing type II errors (Schielzeth, 2010). To evaluate this risk, we computed variance inflation factors (VIF) associated to each term retained in the best model with R package ‘car’ (Fox & Weisberg 2011).

### Statistical analyses of the joint defoliation effects on female fecundity and growth

Our objective herein was to focus on the two variables (growth and female fecundity) responding to defoliation (see results), and to investigate how the relationship between these two variables varied with defoliation. We first compared the effects of defoliation on female fecundity vs growth after centring and normalizing fecundity and growth, and by using the best models fitted with equation 3 to estimate the effect of defoliation on these transformed variables.

Then, we investigated the individual correlation between raw relative female fecundity and growth for non-defoliated trees (DEF=0) versus defoliated trees (DEF>0). Note that a part of these correlations may be due to variation in size and/or competition among individuals. Moreover, they do not account for the quantitative nature of DEF. To overcome these limitations, we further investigated the trade-off between growth and female fecundity using the following linear model:

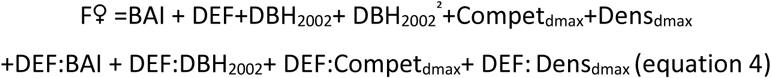

where BAI and the interaction between BAI and DEF are added to the model described by equation (3) above. A quadratic effect of DBH_2002_ was also included.

## Results

### Patterns of covariation of defoliation, tree size and competition

Recurrent crown defoliation was overall limited in the 432 individuals, with 95 trees with a non-null DEF-value (mean=0.36, Table S2). Defoliation increased with tree size; the significant interaction between DBH_2002_ and competition (mediated by Comp19) or density (mediated by Dens20) reflected a stronger effect of size on defoliation as competition increased (Figure S5).

### Inter-individual variations in relative fecundities and growth

The distributions of relative female and male individual fecundities (F♀ and F♂) estimated by MEMM were strongly L-shaped (Figure 3A). Female fecundities varied from 0.03 to 32.44 (median=0.42, mean= 1, sd= 2.78), while male fecundities varied from 0.17 to 21.16 (median= 0.48, mean =1, sd= 1.86). By comparison, the distribution of growth values was less L-shaped than those of fecundity (Figure 3B). In the data set of 432 adult trees, where cumulated growth from 2002 to 2012 was estimated through inventory data, radial growth varied from 0 to 4.4 cm (median=0.45, mean= 0.60, sd= 0.62), while BAI varied from 0 to 581.22 cm^2^ (median=23.98, mean= 61.58, sd= 86.87).

**Figure 3:**
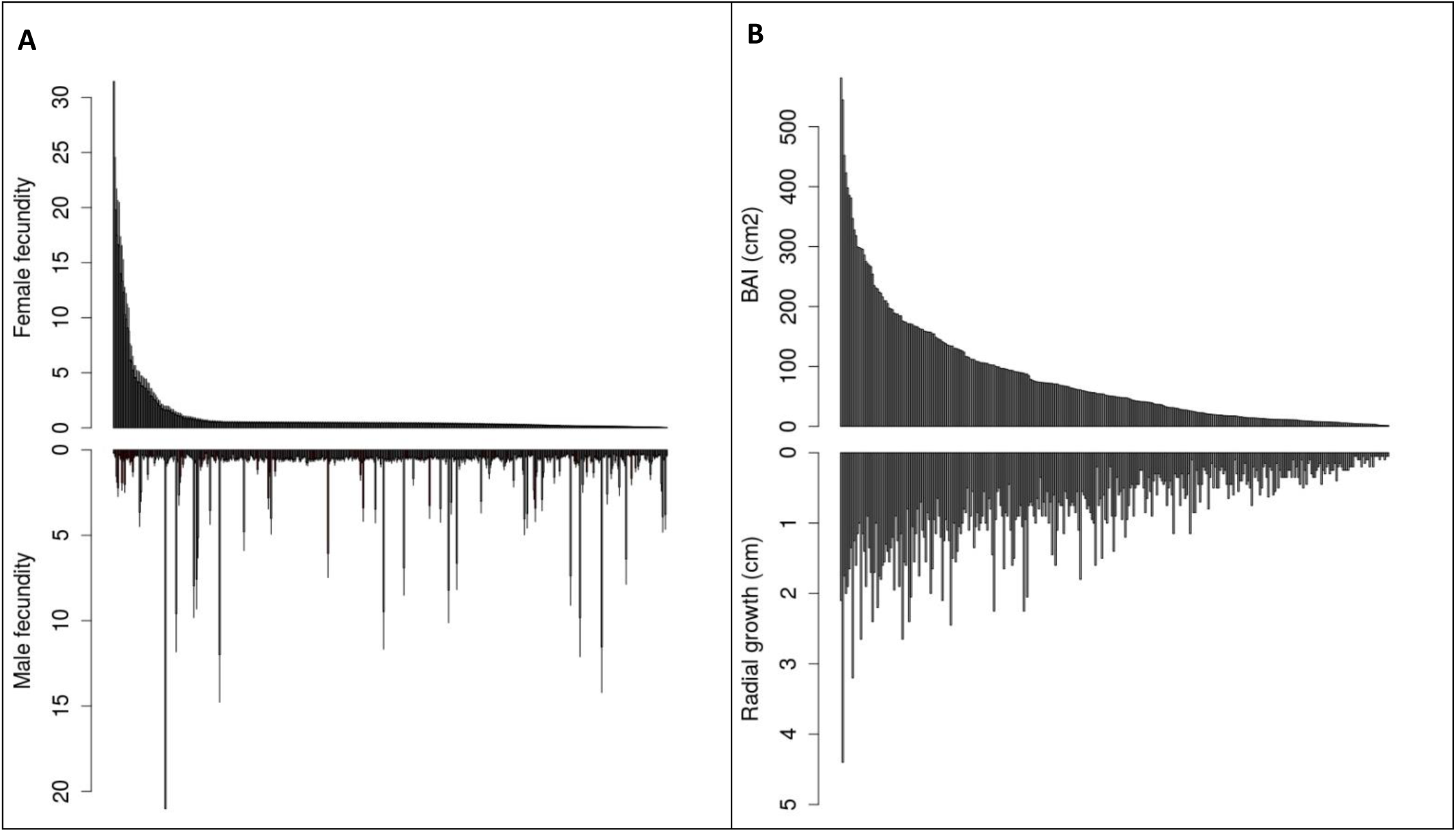
Distribution of individual (A) relative female (top) and male (bottom) fecundities estimated with MEMM, and (B) absolute growth estimated by BAI (top) or radial growth (bottom) for the 432 adult trees. Parents on the x-axis are ranked in decreasing order of female fecundity (A) or BAI (B).

In the subset of 90 cored trees, where cumulated growth from 2002 to 2012 was estimated through ring width data, radial growth varied from 0.17 to 2.70 cm (median=0.97, mean= 1.03, sd= 0.57), while BAI varied from 7.8 to 805.89 cm^2^ (median=126.30, mean= 180.07, sd=172.7). Moreover, for these 90 cored trees, the correlation between inventory-based and ring-width-based radial growth was 0.84 (p-value<0.001), while the correlation between inventory-based and ring-width-based BAI was 0.68 (p-value<0.001). The lower correlation for BAI values was due to the largest trees, for which inventory data generally underestimated growth (Figure S6).

### Ecological drivers of fecundities and growth

Defoliation, size and competition overall explained a significant part of the variation in growth (57%) and female fecundity (12%), while competition alone was found to marginally explain a small part of the variation in male fecundity (<1%). In the whole data set of 432 individuals, the most parsimonious model showed that female fecundity (F♀) significantly decreased with defoliation and competition (mediated by Compet10), while it increased with DBH_2002_ and density (mediated by Dens10; Table 1A). Moreover, the interaction between DEF and DBH_2002_ was significant, reflecting a weaker negative effect of defoliation on female fecundity as tree size increased (Figure 4A). By contrast, male fecundity (F♂) was only marginally (and negatively) affected by competition (mediated by Dens5, Table 1B). Finally, growth (as measured by BAI) significantly decreased with defoliation and competition (mediated by Compet7), and increased with DBH_2002_ and density (mediated by Dens10; Table 1C). By contrast with female fecundity, no interactions between defoliation and size were detected on growth. For all fitted models, variance inflation factors (VIF in Table 1) were all below 10, ruling out any serious multicollinearity issue. Diagnostic plots confirmed the quality of the fitted models (Figure S7).

**Table 1.**
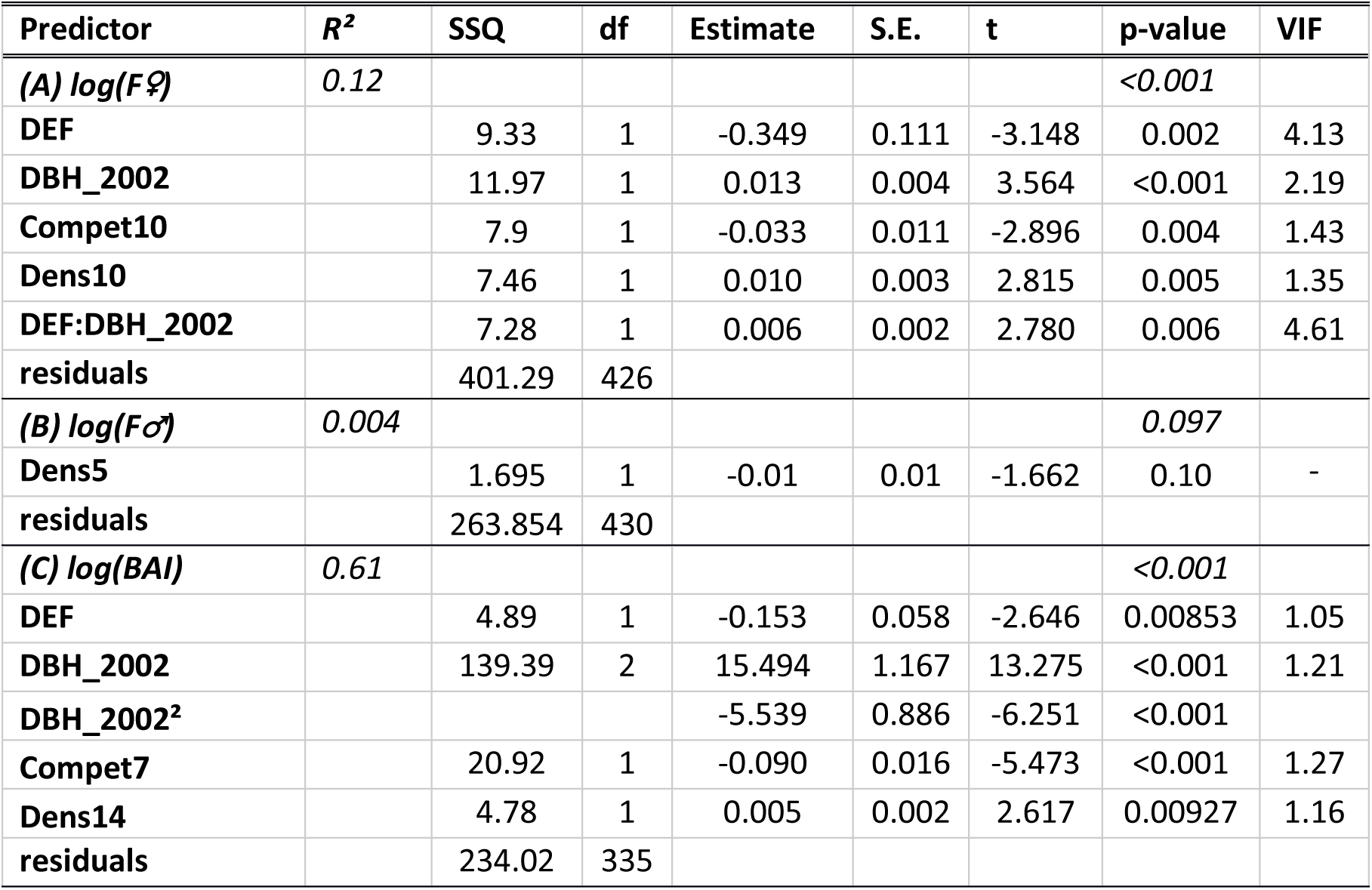
Analysis of variance table for (A) *F♀:* female fecundity (B) *F♂*: male fecundity and (C) BAI: basal area increment, in response to ecological determinants included in equation (3). We show the results of the most parsimonious model: its adjusted R^2^, the type III sum of squares (SSQ) and degree of freedom (df) associated to each term. For each predictor, we give the estimate of its effect, the standard error (S.E.) and associated t and p-value. Variance inflation factors (VIF) were computed with R package CAR. All the response variables were log-transformed. Results are based on the whole data set of 432 individuals for *F♀* and *F♂*, and on the 341 with non-null BAI for BAI.

**Figure 4:**
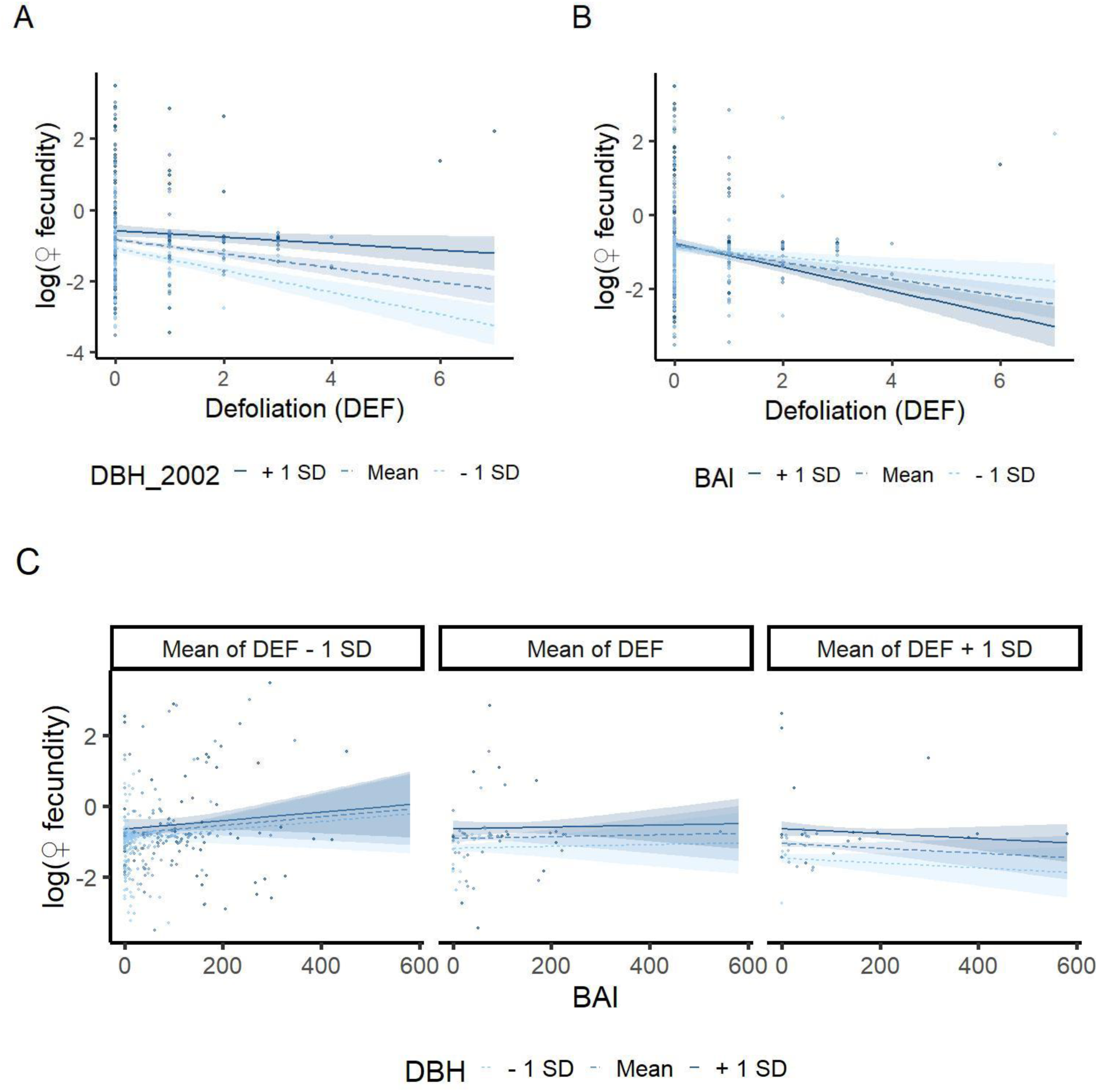
Interaction plots for (A) DEF and DBH_2002_ effects on female fecundity (B) DEF and BAI effects on female fecundity, (C) BAI, DBH_2002_ and DEF effects on female fecundity. Regression lines are plotted for three values of each moderator variable, corresponding to +/- 1 standard deviation from the mean. Confidence interval at 80% are shown around each regression line. Points are the observations.

To compare the effect of defoliation on fecundity and growth, we centred and normalized F♀ and BAI, and ran the best models for each response variable. The average decline in response to a one-unit increase in DEF was −0.06 for F♀ (S.E.= 0.10; measured in standard unit of trait) versus −0.10 for BAI (S.E.=0.04).

### Joint defoliation effects on female fecundity and growth

The raw F♀s and BAIs were significantly and positively correlated in the 337 non-defoliated trees (cor_*F♀-BAI-nondef*_ = 0.31, p-value<0.001), but not in the 95 defoliated trees (cor_*F♀-BAI-def*_=0.13, pval=0.2; Figure 5).

**Figure 5:**
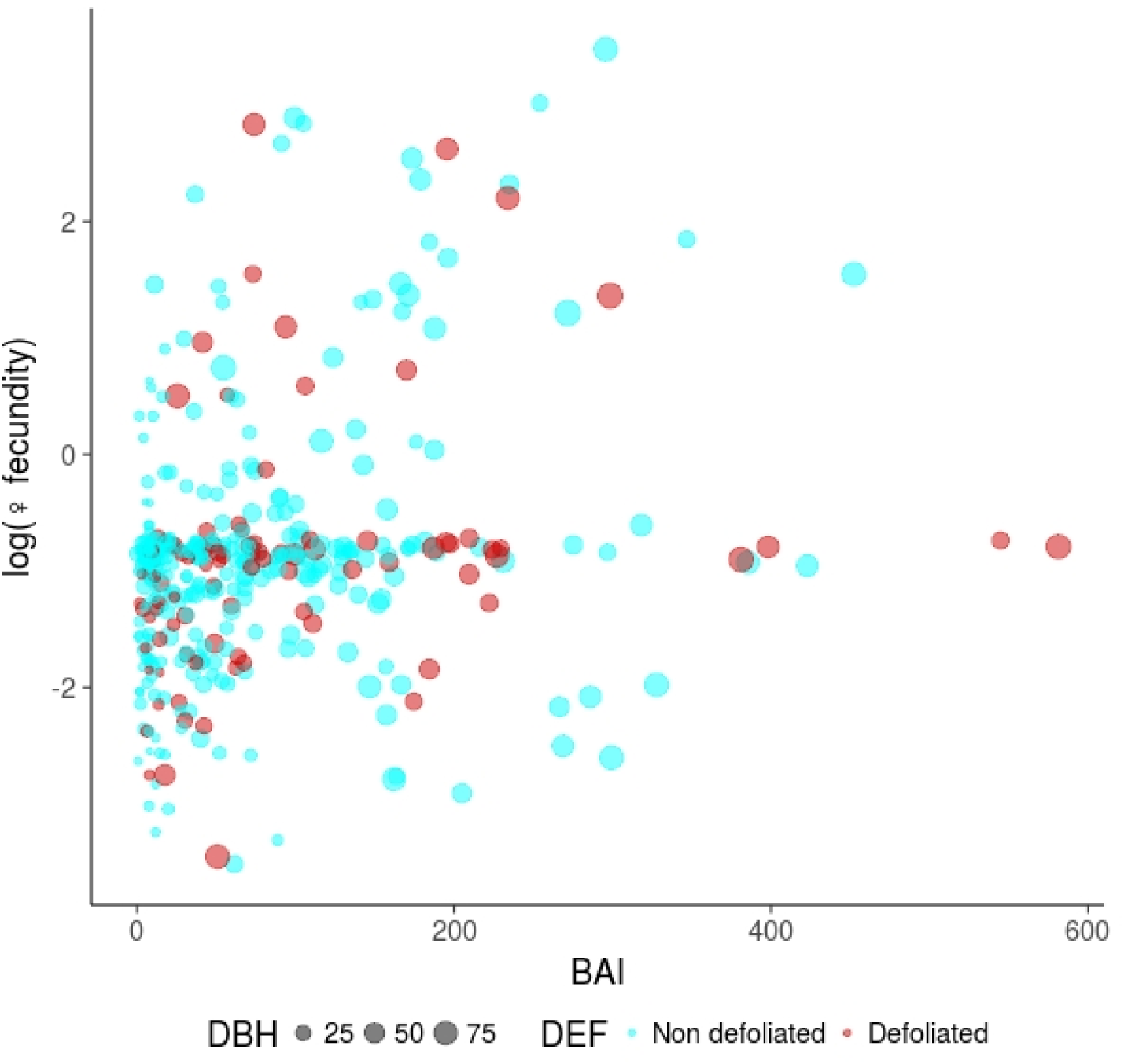
Correlation between growth, measured by the Basal Area Increment (BAI) and female fecundity (F♀), plotted on a log scale. The size of the dots is proportional to tree diameter (DBH_2002_). This is a scatter plot of raw data, and not of model predictions.

The linear model for F♀ including BAI as a predictor (equation 4) allowed us to disentangle the respective effects of defoliation, size and competition on the relationship between female fecundity and growth. In addition to the previous effects, a significant interaction between BAI and defoliation was detected (Table 2): F♀ overall decreased with increasing defoliation, but this decrease was faster and stronger for trees with a higher BAI (Figure 3B). The complex interaction between BAI, DEF and DBH_2002_ on F♀ resulted in a defoliation-dependent trade-off between growth and female fecundity: F♀ of the non-defoliated trees (Figure 3C, left panel) increased with BAI (no trade-off), whereas F♀ of the most defoliated trees (Figure 3C, right panel) decreased with increasing BAI (trade-off). Moreover, F♀ of small trees (Figure S8, left panel) always decreased in response to increasing defoliation, whatever their BAI, whereas the female F♀ of large trees (Figure S8, right panel) could increase in response to increasing DEF, at the expense of reduced BAI. Diagnostic plots confirmed the quality of the fitted models (Figure S9).

**Table 2.** Analysis of variance table for female fecundity, in response to ecological determinants included in equation (4). Results are based on the whole data set of 432 individuals. See Table 1 for legends.

| Predictor | $R^2$ | SSQ | df | Estimate | S.E. | t | P-value | VIF |
| --- | --- | --- | --- | --- | --- | --- | --- | --- |
|  | 0.13 |  |  |  |  |  | <0.001 |  |
| BAI |  | 0.56 | 1 | 0.001 | 0.001 | 0.773 | 0.440 | 2.468 |
| DEF |  | 11.04 | 1 | -0.384 | 0.112 | -3.433 | 0.001 | 4.231 |
| DBH |  | 6.24 | 1 | 0.011 | 0.004 | 2.582 | 0.010 | 3.171 |
| Compet10 |  | 7.49 | 1 | -0.032 | 0.011 | -2.828 | 0.005 | 1.444 |
| Dens10 |  | 6.85 | 1 | 0.009 | 0.003 | 2.704 | 0.007 | 1.354 |
| BAI:DEF |  | 4.09 | 1 | -0.001 | 0.001 | -2.091 | 0.037 | 2.857 |
| DEF:DBH |  | 11.39 | 1 | 0.009 | 0.003 | 3.488 | 0.001 | 6.750 |
| Residuals |  | 397.03 | 424 |  |  |  |  |  |

## Discussion

By investigating the among-individual variation in the impact of stress-induced defoliation on female/male fecundity and wood growth within a beech natural population at the warm, dry margin of the species distribution, this study brings new insights on the response to stress of a major European tree species. We show that crown defoliation was significantly associated to a decrease in wood growth and female fecundity, but not in male fecundity. A trade-off between growth and female fecundity was observed in response to defoliation, suggesting that some large defoliated individuals can maintain significant female fecundity at the expense of reduced growth. The consequences of these results on short-term evolutionary dynamics of the studied population are discussed.

### The response of wood growth and reproduction to stress-induced crown defoliation

The coordination between increasing crown defoliation and decreasing wood growth observed in this study is consistent with the temporal sequence of ecophysiological processes involved in tree response to water stress and late frosts. During summer, low precipitation and high evaporative demand due to high temperatures and vapor-pressure increase water stress, which leads trees to close stomata, in order to reduce transpiration and protect the integrity of the hydraulic system by maintaining water potentials above irreversible embolism thresholds. Drought also directly impacts wood growth by limiting cell division and elongation of wood cells due to carbon limitation (Lempereur et al., 2015). Post-drought stomatal closure can prolong the decrease in photosynthesis and potentially affect carbon storage (Bréda et al., 2006), which may lead to a decrease in radial growth in subsequent years. Under severe drought, some branches can experience hydraulic failure or undergo carbon starvation, which leads to leaf fall. Leaf fall can then in turn have a negative effect on radial growth, first by decreasing photosynthesis and thus carbon availability in the years following defoliation. Secondly, leaf fall can induce allocation shifts that reduce the priority of growth relative to other sinks such as reserves storage, as observed in black oak (Wiley et al., 2017). On the other hand, when late frosts damage young leaves, beech trees can reflush, i.e. produce another cohort of leaves (Menzel, Helm, & Zang, 2015), at least for some parts of the crown. However, the time required to reflush leads to a shorter growing season, which directly reduces wood growth. For these non-mutually exclusive reasons, related to either carbon-, sink-, or temporal limitation of growth, a negative effect of crown defoliation on growth is often observed, especially on beech (Delaporte et al., 2016, this study).

Although seed production is recognized as being resource-limited in plants (Lloyd & Bawa, 1984), the ecophysiological processes involved in the response of tree sexual reproduction to physiological stresses are less well characterized than those involved in wood growth response. The negative effect of crown defoliation on female fecundity observed in this study is consistent with the expected decrease in photosynthesis and thus in carbon availability induced by leaf fall. Moreover, this result combined with the absence of crown defoliation effect on male fecundity is also consistent with the expected higher resource-limitation of female fecundity (costly nut-seeds) as compared to male fecundity in beech (Lloyd & Bawa, 1984; Obeso, 1988). This second expectation is also supported by the marked increase in female fecundity with increasing tree size and decreasing competition while male fecundity was only marginally (and negatively) affected by competition. Moreover, recent studies showed that many tree species use mainly current photosynthates to maturate their fruits, while flowers are produced from old carbon storage (Hoch, Siegwolf, Keel, Körner, & Han, 2013; Ichie et al., 2013). Altogether, our results suggest that beech reproduction is more limited by the carbon resources needed for maturating seeds than for those required for producing flowers. Although nitrogen storage and remobilization is usually a limiting resource for seed production, and particularly masting (Han & Kabeya, 2017), this may not be the case in our study site, where cow grazing could favour nitrogen enrichment.

### Defoliation induced a trade-off between growth and reproduction

Several studies tested the existence of a negative correlation between growth and reproduction at the individual level, as a signature of the possible trade-off between these functions. The key assumption underlying this trade-off is that reproduction is costly and competes with growth for resources (Koenig & Knops, 1998; Obeso, 2002; Thomas, 2011). By contrast, the absence of correlation is usually interpreted as independence between these functions in terms of resource pool (Knops, Koenig, & Carmen, 2007; Obeso, 2002; Pulido et al., 2014). A trade-off between growth and reproduction was already found for beech (Hacket-Pain, Friend, Lageard, & Thomas, 2015; Lebourgeois et al., 2018; Hacket-Pain et al., 2018). More precisely, Hacket-Pain et al. (2017) found for beech that masting years (i.e. high seed production) are negatively correlated with growth and this trade-off is more pronounced during drought years due to resource scarcity.

We found here a negative (respectively positive) correlation between growth and reproduction for defoliated (respectively non-defoliated) trees. Hence, among the defoliated trees, some individuals maintained significant female fecundity at the expense of reduced growth, and reciprocally. These results support the general idea that the correlation between reproduction and growth depends on the level of resource (Obeso, 2002; van Noordwijk & de Jong, 1986), a trade-off being present only under limiting resources, i.e. crown defoliation in our case. By contrast, the higher resource level of non-defoliated individuals could allow them to insure reproduction and growth with independent resource pool, as it was found also for *Fagus* genus (Yasumura, Hikosaka, & Hirose, 2006). Moreover, the detailed analysis of the interactions between defoliation, size and growth on female fecundity showed that those defoliated trees maintaining high female fecundity were the largest ones, suggesting that crown defoliation could shift the allocation of carbon to reproduction above a given tree size. Besides the literature on forest seed orchards and fruit trees orchards, one of the rare studies supporting such hypothesis is that of Wiley, Casper & Helliker (2017), who experimentally defoliated black oak, a tree species which maturates its acorns over two years. Recovery following defoliation was shown to involve substantial allocation shifts, with carbohydrate storage and already initiated reproduction cycles (i.e. maturation of 2-year acorn) being favored relative to growth and new reproductive cycles (i.e. flowering and production of new 1-year acorn).

The positive correlation between growth and reproduction for non-defoliated trees may also indicate an effect of the unobserved level of resource, which probably varies among individuals. More generally, elucidating the causal relationships between defoliation as an impact of stress, the (non-observed) level of resource, growth and reproduction would deserve further investigations, accounting for the complex multivariate relationships among the interrelated variables mapped on Figure 1. This could be achieved using for instance path analyses (Shipley, 2016) or other Bayesian tools introducing the level of resource as a latent variable (e.g., Journé et al., submitted). The use of such approaches in this study was however hampered by two main limitations. First, resource allocation between two compartments are difficult to handle in path analyses, and a reciprocal relationship between growth and reproduction such as depicted by the red double arrow on Figure 1 cannot be specified (Shipley, 2016). The solution to this problem usually consists in accounting for the time dimension, focusing on among-year lagged effects between annual variables (e.g. Hacket-Pain et al., 2018, Journé et al., submitted). However, this solution was intractable in this study, where growth and reproduction were measured as integrated values over the period 2002-2012. The second limitation stems from the weak ability of variance-covariance based methods such as path analyses to deal with non-normality. Whereas deviations from a Gaussian distribution are not necessarily crucial for predictor variables in a linear model, they cannot be handled in path analyses simultaneously with latent variable to our knowledge (Lefcheck, 2016).

### Long-term consequences for population adaptive response to stress

In the studied population, some large, defoliated individuals maintained a high female fecundity under stressful conditions, at the expense of reduced growth. Moreover, as male fecundity was insensitive to crown defoliation, the less competed defoliated trees also contribute to reproduction through male function. Hence, we can reject the hypothesis H1 that the relationship between reproduction and growth does not change with increasing crown defoliation. Note however that the alternative hypothesis H2 (defoliation or stresses act like a cue stimulating reproductive performances at the expense of reduced growth) was only supported for some individuals. This response to stress could have major consequences for the short-term evolutionary dynamics of the population. Indeed, assuming that at least some of the traits underlying vulnerability to stresses are under genetic control, we showed here that the most vulnerable individuals (those that are the most impacted by stress) still contribute to regeneration, which could lead the population to evolve traits compromising its adaptation to stress. By contrast, if the defoliated individuals would decrease simultaneously their growth and reproduction (hypothesis H1), their potentially non-adapted genotypes could be purged more efficiently.

Deriving demo-genetic scenarios for the population adaptive response to stress and testing for a reproduction load would however require further investigations. First, the observed inter-individual variation in the level of defoliation is probably shaped in part by genetic variation but also by microenvironment variation and ontogeny, since the largest and most competed individuals were more susceptible to defoliation. Hence, the importance of genetic factors driving the level of defoliation remains to be characterised, in order to better decipher the intra-individual variation in the vulnerability to stress from that of the stress exposure, and to investigate possible evolutionary changes (Hamanishi & Campbell, 2011). Second, our MEMM-based estimates of fecundity have the advantage to integrate the whole regeneration processes and not just seed/pollen production. This is important as post-dispersal processes and recruitment patterns may compensate the decline in seed production in populations under stressful conditions, as suggested for drought by Barbeta et al. (2011). However, they also have the drawbacks to be relative, and to convey no information on the absolute contribution of defoliated or non-defoliated individuals to the regeneration, and thus on the demographic impact of defoliation.

Finally, investigating the population adaptive response to stress would ideally require accounting for the physiological mechanisms involved. For instance, when dealing with drought-induced defoliation, we would need to consider the two main ecological strategies widely acknowledged in plants for drought response: 1) the water economy strategy, where plants maintain low growth rates and low rates of gas exchange during droughts, and 2) the water uptake strategy, where plants have a more rapid instant growth through higher rates of gas exchange when water is available, typically spring in Mediterranean climate, allowing them to complete important biological functions before drought onset (Arntz & Delph, 2001). These two strategies rely on different combinations of physiological, morphological or phenological trait values (in particular those related to hydraulics and carbon-storage). Bontemps et al. (2017) demonstrated the co-existence of these two strategies in a drought-prone population of beech and showed a higher reproductive output of the water uptake strategy. In this context, defoliation could be one of the traits involved in the water uptake strategy, allowing the maintenance of the water balance after drought onset. Indeed, if defoliated trees are characterized by higher xylem vulnerability but also higher hydraulic conductivity, the more a tree is efficient for transpiration and photosynthesis, the more it is vulnerable to drought (Cochard, Lemoine, & Dreyer, 1999).

## Data accessibility

The data set analysed in this preprint is available online under the zenodo platform (DOI: 10.5281/zenodo.3516305). More detailed data (genotypes) and R scripts for statistical analyses are available from the corresponding author.

## Supplementary material

The MEMM software for estimating female and male fecundities is available at: https://informatique-mia.inra.fr/biosp/memm.

Supplementary materials (Figures and Tables) for this preprint are available on bioRxiv (doi: https://doi.org/10.1101/474874).

## Acknowledgements

We are grateful to our colleagues Francois Lefèvre and Etienne K. Klein, as well as to our PCIEcology editor Georges Kunstler and three anonymous reviewers for discussions and comments on previous version of this manuscript. We thank Nicolas Mariotte INRA URFM Avignon for wood core sampling and Jean Thevenet, INRA UEFM, Avignon for sample management. The study was partly funded by the EU ERA-NET BiodivERsA projects TIPTREE (BiodivERsA2-2012-15), and the ANR project MeCC (ANR-13-ADAP-0006). Version 4 of this preprint has been peer-reviewed and recommended by Peer Community In Ecology (https://doi.org/10.24072/pci.ecology.100033).

## Conflict of interest disclosure

The authors of this preprint declare that they have no financial conflict of interest with the content of this article. SOM is a recommender for PCIEcology.

## Notes

#### Summary of Updates

Version 4 of this preprint has been peer-reviewed and recommended by Peer Community In Ecology (https://doi.org/10.24072/pci.ecology.100033).

https://zenodo.org/record/3516305#.Xb269tXjJPY

## References

Adams, H. D., Zeppel, M. J. B., Anderegg, W. R. L., Hartmann, H., Landhäusser, S. M., Tissue, D. T., … McDowell, N. G. (2017). A multi-species synthesis of physiological mechanisms in drought-induced tree mortality. Nature Ecology and Evolution, 1(9), 1285–1291. doi:10.1038/s41559-017-0248-x

Allen, C. D., Macalady, A. K., Chenchouni, H., Bachelet, D., McDowell, N., Vennetier, M., … Cobb, N. (2010). A global overview of drought and heat-induced tree mortality reveals emerging climate change risks for forests. Forest Ecology and Management, 259(4), 660–684. doi:10.1016/j.foreco.2009.09.001

Anderegg, W. R. L., Kane, J. M., & Anderegg, L. D. L. (2013). Consequences of widespread tree mortality triggered by drought and temperature stress. Nature Climate Change, 3(1), 30–36. doi:10.1038/nclimate1635

Barbeta, A., Peñuelas, J., Ogaya, R., & Jump, A. S. (2011). Reduced tree health and seedling production in fragmented Fagus sylvatica forest patches in the Montseny Mountains (NE Spain). Forest Ecology and Management, 261(11), 2029–2037. doi:10.1016/j.foreco.2011.02.029

Bigler, C., & Bugmann, H. (2018). Climate-induced shifts in leaf unfolding and frost risk of European trees and shrubs. Scientific Reports, 8(1), 1–10. doi:10.1038/s41598-018-27893-1

Bonnet-Masimbert, M., & Webber, J. E. (2012). From flower induction to seed production in forest tree orchards. Tree Physiology, 15(7–8), 419–426. doi:10.1093/treephys/15.7-8.419

Bontemps, A., Davi, H., Lefèvre, F., Rozenberg, P., & Oddou-Muratorio, S. (2017). How do functional traits syndromes covary with growth and reproductive performance in a water-stressed population of Fagus sylvatica? Oikos, 126(10), 1472–1483. doi:10.1111/oik.04156

Bréda, N., Huc, R., Granier, A., & Dreyer, E. (2006). Temperate forest trees and stands under severe drought: a review of ecophysiological responses, adaptation processes and long-term consequences. Annals of Forest Science, 63(6), 625–644. doi:10.1051/forest:2006042

Burczyk, J., Adams, W. T., Birkes, D. S., & Chybicki, I. J. (2006). Using genetic markers to directly estimate gene flow and reproductive success parameters in plants on the basis of naturally regenerated seedlings. Genetics, 173(1), 363–372. doi:10.1534/genetics.105.046805

Bykova, O., Limousin, J.-M., Ourcival, J.-M., & Chuine, I. (2018). Water deficit disrupts male gametophyte development in Quercus ilex. Plant Biology, 20(3), 450–455. doi:10.1111/plb.12692

Bykova, O, Chuine, I., Morin, X., & Higgins, S. I. (2012). Temperature dependence of the reproduction niche and its relevance for plant species distributions. Journal of Biogeography, 39(12), 2191–2200. doi:10.1111/j.1365-2699.2012.02764.x

Camarero, J. J., Gazol, A., Sangüesa-Barreda, G., Oliva, J., & Vicente-Serrano, S. M. (2015). To die or not to die: Early warnings of tree dieback in response to a severe drought. Journal of Ecology, 103(1), 44–57. doi:10.1111/1365-2745.12295

Charrier, G., Ngao, J., Saudreau, M., & Améglio, T. (2015). Effects of environmental factors and management practices on microclimate, winter physiology, and frost resistance in trees. Frontiers in Plant Science, 6(April), 1–18. doi:10.3389/fpls.2015.00259

Delaporte, A., Bazot, S., & Damesin, C. (2016). Reduced stem growth, but no reserve depletion or hydraulic impairment in beech suffering from long-term decline. Trees – Structure and Function, 30(1), 265–279. doi:10.1007/s00468-015-1299-8

Dobbertin, M. (2005). Tree growth as indicator of tree vitality and of tree reaction to environmental stress: a review. European Journal of Forest Research, 124(4), 319–333. doi:10.1007/s10342-005-0085-3

Flores-Rentería, L., Whipple, A. V., Benally, G. J., Patterson, A., Canyon, B., & Gehring, C. A. (2018). Higher Temperature at Lower Elevation Sites Fails to Promote Acclimation or Adaptation to Heat Stress During Pollen Germination. Frontiers in Plant Science, 9, 1–14. doi:10.3389/fpls.2018.00536

Galiano, L., Martínez-Vilalta, J., & Lloret, F. (2011). Carbon reserves and canopy defoliation determine the recovery of Scots pine 4 yr after a drought episode. New Phytologist, 190(3), 750–759. doi:10.1111/j.1469-8137.2010.03628.x

Goto, S., Shimatani, K., Yoshimaru, H., & Takahashi, Y. (2006). Fat-tailed gene flow in the dioecious canopy tree species Fraxinus mandshurica var. japonica revealed by microsatellites. Molecular Ecology, 15(10), 2985–2996. doi:10.1111/j.1365-294X.2006.02976.x

Hacket-Pain, A. J., Ascoli, D., Vacchiano, G., Biondi, F., Cavin, L., Conedera, M., … Zang, C. S. (2018). Climatically controlled reproduction drives interannual growth variability in a temperate tree species. Ecology Letters, 21(12), 1833–1844. doi:10.1111/ele.13158

Hacket-Pain, A. J., Lageard, J. G. A., & Thomas, P. A. (2017). Drought and reproductive effort interact to control growth of a temperate broadleaved tree species (Fagus sylvatica). Tree Physiology, 37(6), 744–754. doi:10.1093/treephys/tpx025

Hamanishi, E. T., & Campbell, M. M. (2011). Genome-wide responses to drought in forest trees. Forestry, 84(3), 273–283. doi:10.1093/forestry/cpr012

Hampe, A., & Petit, R. J. (2005). Conserving biodiversity under climate change: The rear edge matters. Ecology Letters, 8(5), 461–467. doi:10.1111/j.1461-0248.2005.00739.x

Han, Q., & Kabeya, D. (2017). Recent developments in understanding mast seeding in relation to dynamics of carbon and nitrogen resources in temperate trees. Ecological Research, 32(6), 771–778. doi:10.1007/s11284-017-1494-8

Hedhly, A., Hormaza, J. I., & Herrero, M. (2009). Global warming and sexual plant reproduction. Trends in Plant Science, 14(1), 30–36. doi:10.1016/j.tplants.2008.11.001

Hoch, G., Siegwolf, R. T. W., Keel, S. G., Körner, C., & Han, Q. (2013). Fruit production in three masting tree species does not rely on stored carbon reserves. Oecologia, 171(3), 653–662. doi:10.1007/s00442-012-2579-2

Ichie, T., Igarashi, S., Yoshida, S., Kenzo, T., Masaki, T., & Tayasu, I. (2013). Are stored carbohydrates necessary for seed production in temperate deciduous trees? Journal of Ecology, 101(2), 525–531. doi:10.1111/1365-2745.12038

Journé, V., Papaïx, J., Walker, E., Courbet, F., Lefevre, F., Oddou-Muratorio, S., & Davi, H. (submitted). A hierarchical Bayesian resource model to investigate trade-offs between growth and reproduction in a long-lived plant.

Karimi, F., Igata, M., Baba, T., Noma, S., Mizuta, D., Gook Kim, J., & Ban, T. (2017). Summer Pruning Differentiates Vegetative Buds to Flower Buds in the Rabbiteye Blueberry (*Vaccinium virgatum* Ait.). The Horticulture Journal, 86(3), 300–304. doi:10.2503/hortj.MI-158

Lee, T. D. (1988). Patterns of fruit and seed production. In J. Lovett-Doust & L. Lovett-Doust (Eds.), Plant reproductive ecology: patterns and strategies (Oxford Uni). New York.

Lefcheck, J. S. (2016). piecewiseSEM: Piecewise structural equation modelling in r for ecology, evolution, and systematics. Methods in Ecology and Evolution, 7(5), 573–579. doi:10.1111/2041-210X.12512

Lempereur, M., Martin-StPaul, N. K., Damesin, C., Joffre, R., Ourcival, J.-M., Rocheteau, A., & Rambal, S. (2015). Growth duration is a better predictor of stem increment than carbon supply in a Mediterranean oak forest: implications for assessing forest productivity under climate change. New Phytologist, 207(3), 579–590. doi:10.1111/nph.13400

Lloyd, D. G., & Bawa, K. S. (1984). Modification of the Gender of Seed Plants in Varying Conditions. In M. K. Hecht & G. T. Prance (Eds.), Evolutionary Biology (pp. 255–338). Boston, MA: Routledge. doi:10.4324/9781315128634-1

McDowell, N. G., Beerling, D. J., Breshears, D. D., Fisher, R. A., Raffa, K. F., & Stitt, M. (2011). The interdependence of mechanisms underlying climate-driven vegetation mortality. Trends in Ecology and Evolution, 26(10), 523–532. doi:10.1016/j.tree.2011.06.003

Meilan, R. (1997). Floral induction in woody angiosperms. New Forests, 14, 179–202.

Menzel, A., Helm, R., & Zang, C. (2015). Patterns of late spring frost leaf damage and recovery in a European beech (Fagus sylvatica L.) stand in south-eastern Germany based on repeated digital photographs. Frontiers in Plant Science, 6, 1–13. doi:10.3389/fpls.2015.00110

Moran, E. V., & Clark, J. S. (2011). Estimating seed and pollen movement in a monoecious plant: A hierarchical Bayesian approach integrating genetic and ecological data. Molecular Ecology, 20(6), 1248–1262. doi:10.1111/j.1365-294X.2011.05019.x

Obeso, J. R. (1988). The costs of reproduction in plants. New Phytologist, 155:, 321–348. doi:10.1046/j.1469-8137.2002.00477.x

Oddou-Muratorio, S., Gauzere, J., Bontemps, A., Rey, J. F., & Klein, E. K. (2018). Tree, sex and size: Ecological determinants of male vs. female fecundity in three Fagus sylvatica stands. Molecular Ecology, 27(15), 3131–3145. doi:10.1111/mec.14770

Oddou-Muratorio, S., & Klein, E. K. (2008). Comparing direct vs. indirect estimates of gene flow within a population of a scattered tree species. Molecular Ecology, 17(11), 2743–2754. doi:10.1111/j.1365-294X.2008.03783.x

Peñuelas, J., & Boada, M. (2003). A global change-induced biome shift in the Montseny mountains (NE Spain). Global Change Biology, 9(2), 131–140. doi:10.1046/j.1365-2486.2003.00566.x

Pérez-Ramos, I. M., Ourcival, J. M., Limousin, J. M., & Rambal, S. (2010). Mast seeding under increasing drought: results from a long-term data set and from a rainfall exclusion experiment. Ecology, 91(10), 3057–3068. doi:10.1890/09-2313.1

Petit-Cailleux, C., Davi, H., Lefevre, F., Garrigue, J., Magdalou, J.-A., Hurson, C., … Oddou-Muratorio, S. (under review). Combining statistical and mechanistic models to unravel the drivers of mortality within a rear-edge beech population. PCIEcology.

Pulido, F., Moreno, G., Garcia, E., Obrador, J. J., Bonal, R., & Diaz, M. (2014). Resource manipulation reveals flexible allocation rules to growth and reproduction in a Mediterranean evergreen oak. Journal of Plant Ecology, 7(1), 77–85. doi:10.1093/jpe/rtt017

Quézel, P., & Médail, F. (2003). Écologie et biogéographie des forêts du bassin méditerranéen. Elsevier.

Sánchez-Humanes, B., & Espelta, J. M. (2011). Increased drought reduces acorn production in Quercus ilex coppices: Thinning mitigates this effect but only in the short term. Forestry, 84(1), 73–82. doi:10.1093/forestry/cpq045

Schielzeth, H. (2010). Simple means to improve the interpretability of regression coefficients. Methods in Ecology and Evolution, 1(2), 103–113. doi:10.1111/j.2041-210X.2010.00012.x

Sherry, R. A., Zhou, X., Gu, S., Arnone, J. A., Schimel, D. S., Verburg, P. S., … Luo, Y. (2007). Divergence of reproductive phenology under climate warming. Proceedings of the National Academy of Sciences, 104(1), 198–202. doi:10.1073/pnas.0605642104

Shipley, B. (2016). Cause and Correlation in Biology. Cause and Correlation in Biology. doi:10.1017/cbo9781139979573

Vacchiano, G., Hacket-Pain, A., Turco, M., Motta, R., Maringer, J., Conedera, M., … Ascoli, D. (2017). Spatial patterns and broad-scale weather cues of beech mast seeding in Europe. New Phytologist, 215(2), 595–608. doi:10.1111/nph.14600

Wiley, E., Casper, B. B., & Helliker, B. R. (2017). Recovery following defoliation involves shifts in allocation that favour storage and reproduction over radial growth in black oak. Journal of Ecology, 105(2), 412–424. doi:10.1111/1365-2745.12672

Yasumura, Y., Hikosaka, K., & Hirose, T. (2006). Resource allocation to vegetative and reproductive growth in relation to mast seeding in Fagus crenata. Forest Ecology and Management, 229(1–3), 228–233. doi:10.1016/j.foreco.2006.04.003

Zhao, M., & Running, S. W. (2010). Drought-Induced Reduction in Global Terrestrial Net Primary Production from 2000 Through 2009. Science, 329(5994), 940–943. doi:10.1126/science.1192666

Zimmermann, J., Hauck, M., Dulamsuren, C., & Leuschner, C. (2015). Climate Warming-Related Growth Decline Affects Fagus sylvatica, But Not Other Broad-Leaved Tree Species in Central European Mixed Forests. Ecosystems, 18(4), 560–572. doi:10.1007/s10021-015-9849-x

Zinn, K. E., Tunc-Ozdemir, M., & Harper, J. F. (2010). Temperature stress and plant sexual reproduction: Uncovering the weakest links. Journal of Experimental Botany, 61(7), 1959–1968. doi:10.1093/jxb/erq053

