## Supplementary online material for "Crown defoliation decreases reproduction and wood growth in a marginal European beech population"

**Table S1: Genetic data for the adult and offspring population.**

For each SSR marker, N is the total number of scored individuals, %M the percentage of missing data. Na is the number of alleles, AR allelic richness (estimated for a sample of 36 individuals), Ne the effective number of alleles ; He the expected heterozygosity. Exclusion probabilities for maternity (PE-1P) and parentage (PE-PP) are also given with their cumulated values over 21 loci on the last line.

| Marker | N | %M | Na | AR | Ne | He | PE-1P | PE-PP |
|---|---|---|---|---|---|---|---|---|
| **Csolfagus_19** | 956 | 8.8% | 12 | 8.07 | 5.69 | 0.82 | 0.52 | 0.17 |
| **Csolfagus_7** | 726 | 30.7% | 6 | 5.04 | 4.50 | 0.78 | 0.63 | 0.27 |
| **F1_15** | 928 | 11.5% | 18 | 9.83 | 4.77 | 0.79 | 0.56 | 0.18 |
| **Fs3_4** | 821 | 21.7% | 4 | 2.69 | 2.02 | 0.50 | 0.87 | 0.69 |
| **Sfc0007** | 854 | 18.5% | 7 | 5.21 | 4.20 | 0.76 | 0.65 | 0.29 |
| **Sfc1143** | 746 | 28.8% | 10 | 7.05 | 3.17 | 0.68 | 0.70 | 0.31 |
| **Csolfagus_25** | 906 | 13.5% | 6 | 4.73 | 2.17 | 0.54 | 0.85 | 0.53 |
| **Csolfagus_29** | 908 | 13.4% | 5 | 4.04 | 2.68 | 0.63 | 0.79 | 0.45 |
| **Csolfagus_31** | 742 | 29.2% | 11 | 6.65 | 3.17 | 0.68 | 0.72 | 0.36 |
| **Csolfagus_6** | 1025 | 2.2% | 10 | 6.66 | 4.23 | 0.76 | 0.63 | 0.27 |
| **Fi05** | 297 | 71.7% | 7 | 5.36 | 1.99 | 0.50 | 0.86 | 0.54 |
| **Mfc7** | 915 | 12.7% | 7 | 5.76 | 1.98 | 0.49 | 0.86 | 0.50 |
| **sfc061** | 559 | 46.7% | 13 | 9.72 | 5.19 | 0.81 | 0.54 | 0.18 |
| **concat14_A_0** | 689 | 34.3% | 6 | 5.15 | 3.50 | 0.71 | 0.70 | 0.35 |
| **DE576** | 552 | 47.3% | 6 | 4.55 | 3.16 | 0.68 | 0.74 | 0.41 |
| **DUKCT_A_0** | 920 | 12.2% | 5 | 4.97 | 2.66 | 0.62 | 0.78 | 0.41 |
| **DZ447_A_0** | 1019 | 2.8% | 6 | 5.46 | 3.70 | 0.73 | 0.68 | 0.31 |
| **EEU75** | 891 | 15.0% | 9 | 5.26 | 1.91 | 0.48 | 0.88 | 0.55 |
| **EJV8T** | 1001 | 4.5% | 8 | 4.53 | 2.61 | 0.62 | 0.80 | 0.50 |
| **EMILY_A** | 869 | 17.1% | 6 | 5.50 | 3.06 | 0.67 | 0.74 | 0.39 |
| **ERHIBI_A_0** | 718 | 31.5% | 6 | 3.58 | 2.75 | 0.64 | 0.80 | 0.49 |
| **Mean** | 811.5 | 22.6% | 8.0 | 5.7 | 3.29 | 0.66 | | |
| **Cumulated** | | | | | | | 0.001 | $5.8\ 10^{-10}$ |

**Table S2: Size, competition and defoliation variables measured in adult trees.**

| Trait code | Trait | Unit | Value | 432 trees | 90 cored trees |
|---|---|---|---|---|---|
| **$DBH_{2002}$ (cm)** | Diameter in 2002 | cm | ***Mean (sd)*** | 25.7 (18.9) | 39.5 (14.5) |
| | | | ***Median*** | 22.4 | 34.2 |
| | | | ***Min-max*** | 2.5-95.2 | 18.5-95.2 |
| **Compet5** | Competition index within 5 m | - | ***Mean (sd)*** | 3.9 (3.5) | 1.90 (1.1) |
| | | | ***Median*** | 3 | 1.7 |
| | | | ***Min-max*** | 0– 23.1 | 0.03-4.6 |
| **Compet10** | Competition index within 10 m | - | ***Mean (sd)*** | 6.6 (5.0) | 3.7 (1.2) |
| | | | ***Median*** | 5.2 | 3.7 |
| | | | ***Min-max*** | 0.8 – 30.5 | 0.8-6.5 |
| **Compet15** | Competition index within 15 m | - | ***Mean (sd)*** | 8.05 (5.5) | 4.9 (1.2) |
| | | | ***Median*** | 6.5 | 5 |
| | | | ***Min-max*** | 0.6 – 30.8 | 1.9-7.7 |
| **Compet20** | Competition index within 20 m | - | ***Mean (sd)*** | 9.47 (7.0) | 6.9 (6.0) |
| | | | ***Median*** | 7.1 | 5.8 |
| | | | ***Min-max*** | 2.5 – 34.0 | 2.6-33.9 |
| **Dens5** | Nb of neighbors within 5 m | - | ***Mean (sd)*** | 9.7 (5.3) | 8.1 (5.1) |
| | | | ***Median*** | 10 | 7.5 |
| | | | ***Min-max*** | 0-24 | 1-22 |
| **Dens10** | Nb of neighbors within 10 m | - | ***Mean (sd)*** | 34.0 (15.6) | 29.6 (15.3) |
| | | | ***Median*** | 33 | 26.5 |
| | | | ***Min-max*** | 4-70 | 5-65 |
| **Dens15** | Nb of neighbors within 15 m | - | ***Mean (sd)*** | 75.1 (34.1) | 8.1 (5.1) |
| | | | ***Median*** | 72 | 57 |
| | | | ***Min-max*** | 11-166 | 14-124 |
| **Dens20** | Nb of neighbors within 20 m | - | ***Mean (sd)*** | 132.9 (60.3) | 116.9 (55.0) |
| | | | ***Median*** | 135 | 103 |
| | | | ***Min-max*** | 20-263 | 28-219 |
| **DEF** | Defoliation index | - | ***Mean (sd)*** | 0.37 (0.9) | 0.7 (1.0) |
| | | | ***Median*** | 0 | 0 |
| | | | ***Min-max*** | 0-7 | 0 -4 |

**Figure S1 Climate characteristics of the study site.**

**Figure S1A:** Climatic diagram at La Massane representing the sum of monthly precipitations (P, blue barplot) and the average monthly temperature (T, continuous black line). The error bars on the P barplot and the dashed lines around the T continuous line show the confidence interval at 95% of monthly values, based on the variation observed from 1976 to 2015

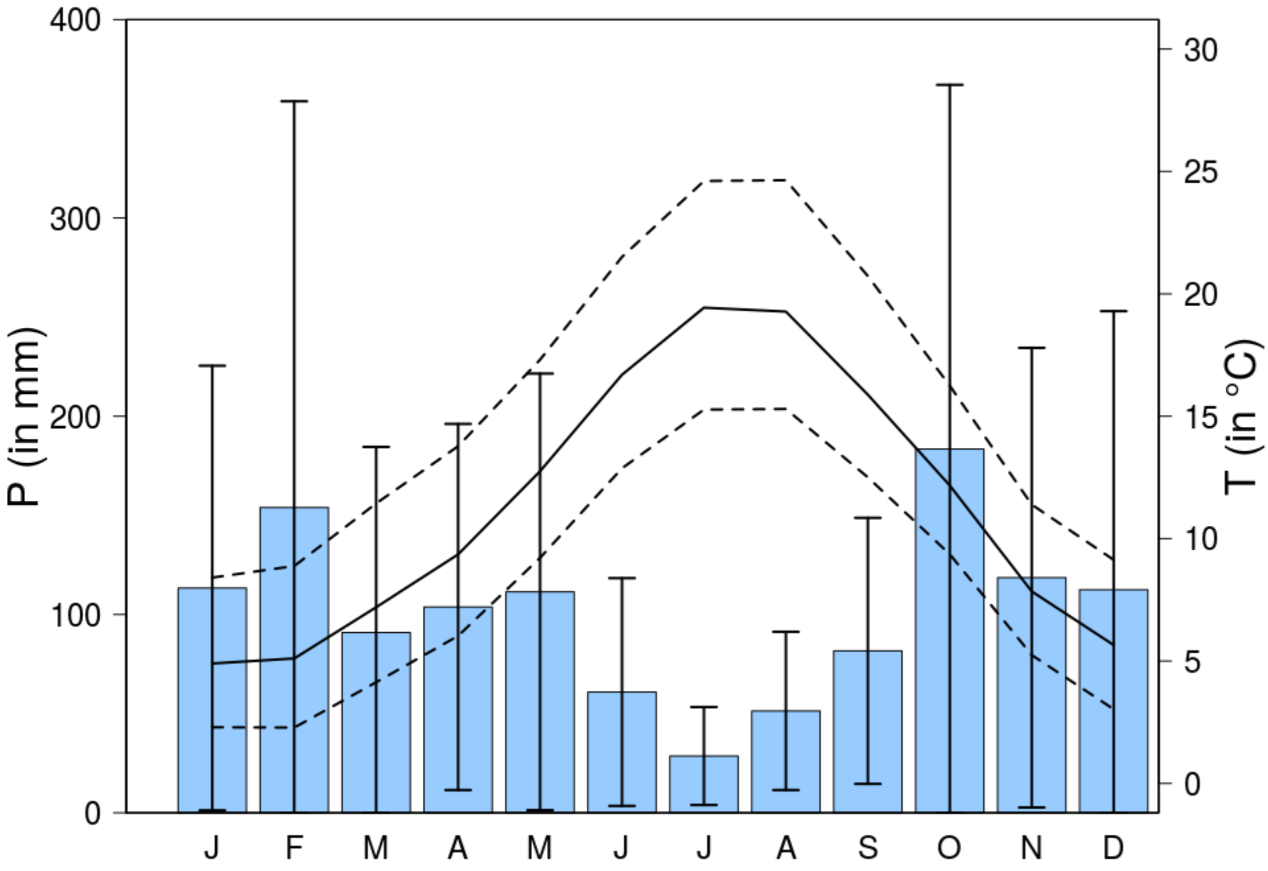

**Figure S1B:** Position of La Massane (as the magenta triangle) on French beech bioclimatic niche (green crosses). Left: presence of beech (green crosses) according to French Inventory data (IFN). We used the meteorological Safran data base for the period 1958-2015 (collected on a 8 km-square grid represented by black empty circles) to draw the bioclimatic niche graph (right), as depicted by mean annual temperature (MAT) and the sum of summer precipitation (PRECsummer). Note that the magenta triangle correspond the the Safran point the closest from La Massane, and not to the climate data monitored on site as in Figure S1.

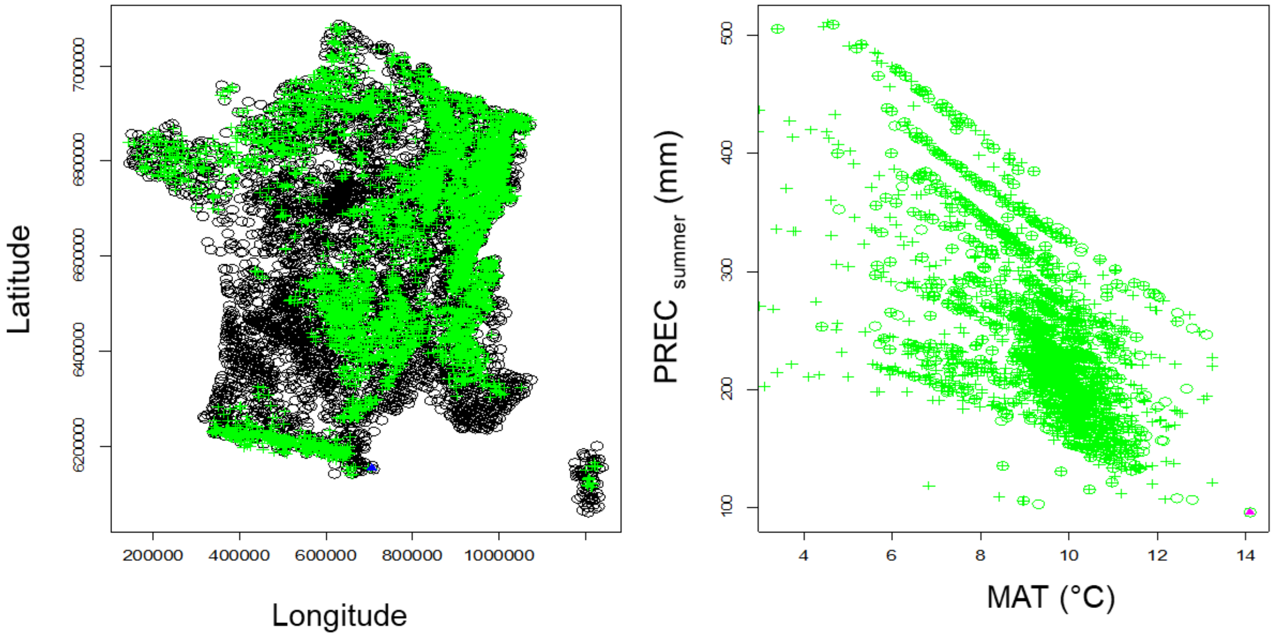

**Figure S2: Patterns of covariation among competition index (the CX's) and density (the DX's) computed in radius a different size (X=1 to 20 m) around each focal beech.**

The left plot shows the variables projection onto the Principal Component Analysis plane define by the two first axis. The right plot shows the pairwise correlations between variables.

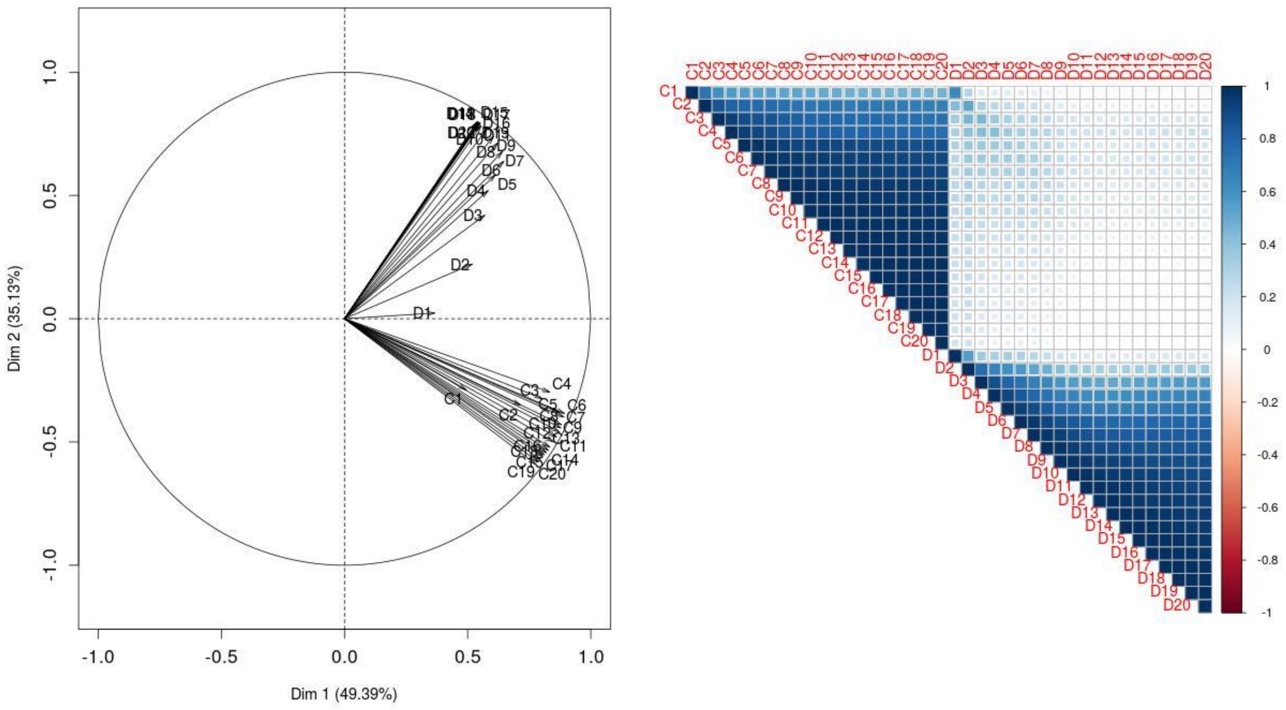

**Figure S3: Sampling design for the 90 cored individuals**
**A. Map of the cored trees per category.** Points represent all the 683 adult alive beeches.

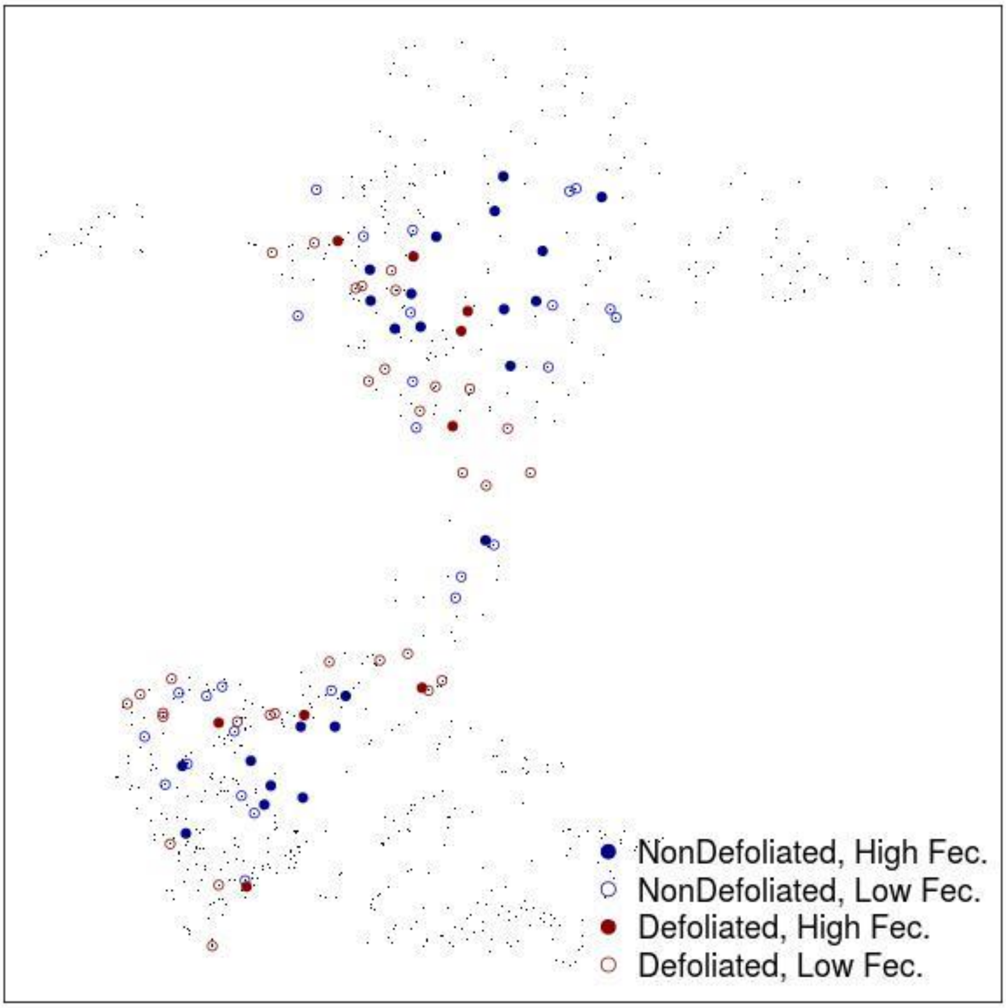

**B. Sampling size, mean (sd) fecundity and defoliation per category**

| **Category** | **# indiv.** | **Female fecundity*** | **Defoliation ** ** |
|---|---|---|---|
| **Non-defoliated, High fecundity** | 23 | 2.1 (2.2) | 0 (0) |
| **Non-defoliated, Low fecundity** | 27 | 0.16 (0.16) | 0 (0) |
| **Defoliated, High fecundity** | 9 | 1.1 (1.6) | 1.4 (0.73) |
| **Defoliated, Low fecundity** | 31 | 0.13 (0.051) | 1.7 (0.98) |

*Being relative, fecundity values have no unit.
** Defoliation is estimated as the sum of annual defoliation scores (0= absence versus 1= presence of dead branches/leaves) over 9 years; so they also have no units.

**Figure S4: Preliminary check of the quality of linear models described by equation (3) and (4).**

A. Distribution of predictor variables (not transformed)
B. Distribution of response variables (before and after log-transformation)
C. Relationship between each predictor and each response variable of the model described by equation (3) and (4).

A.

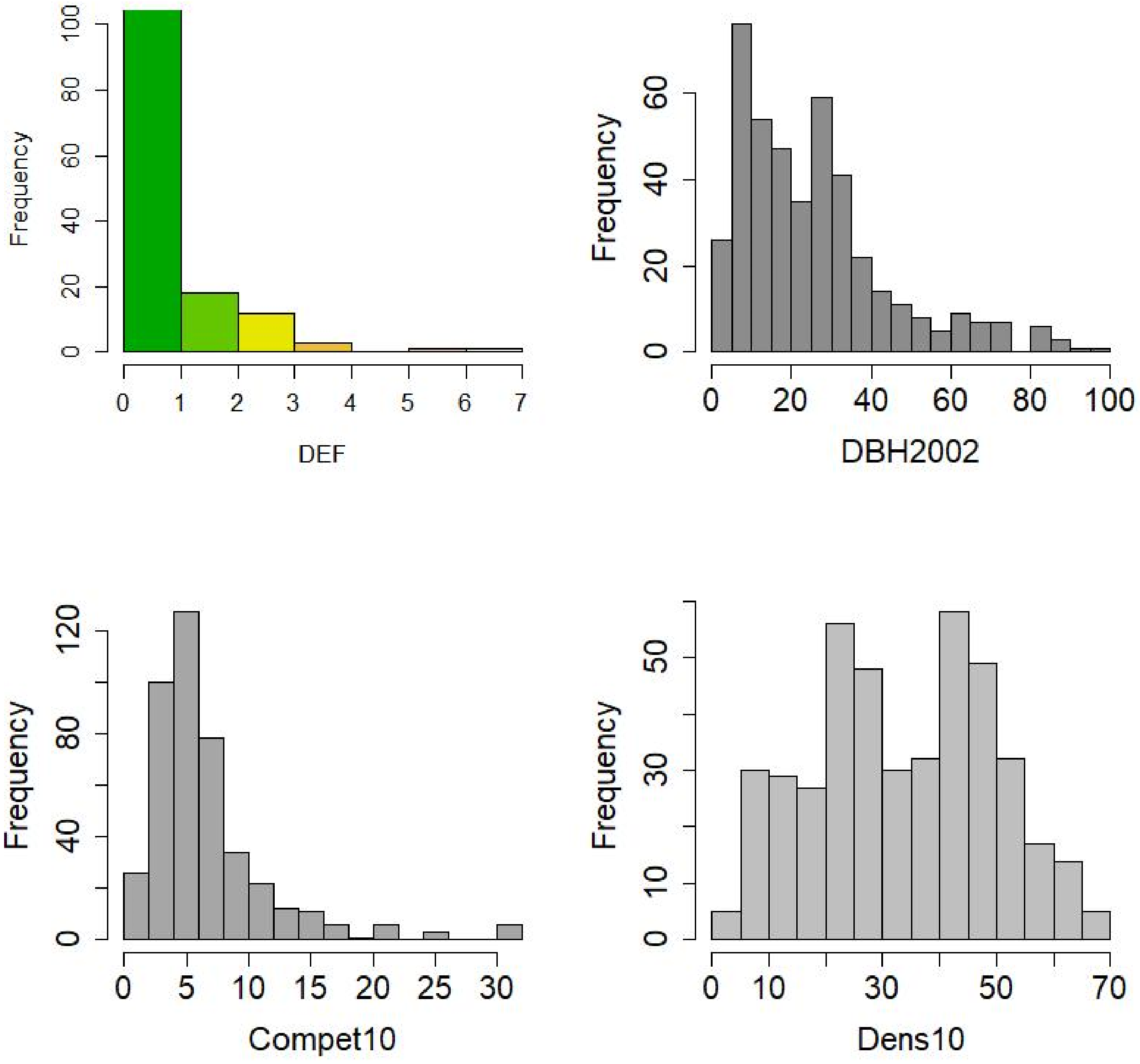

B

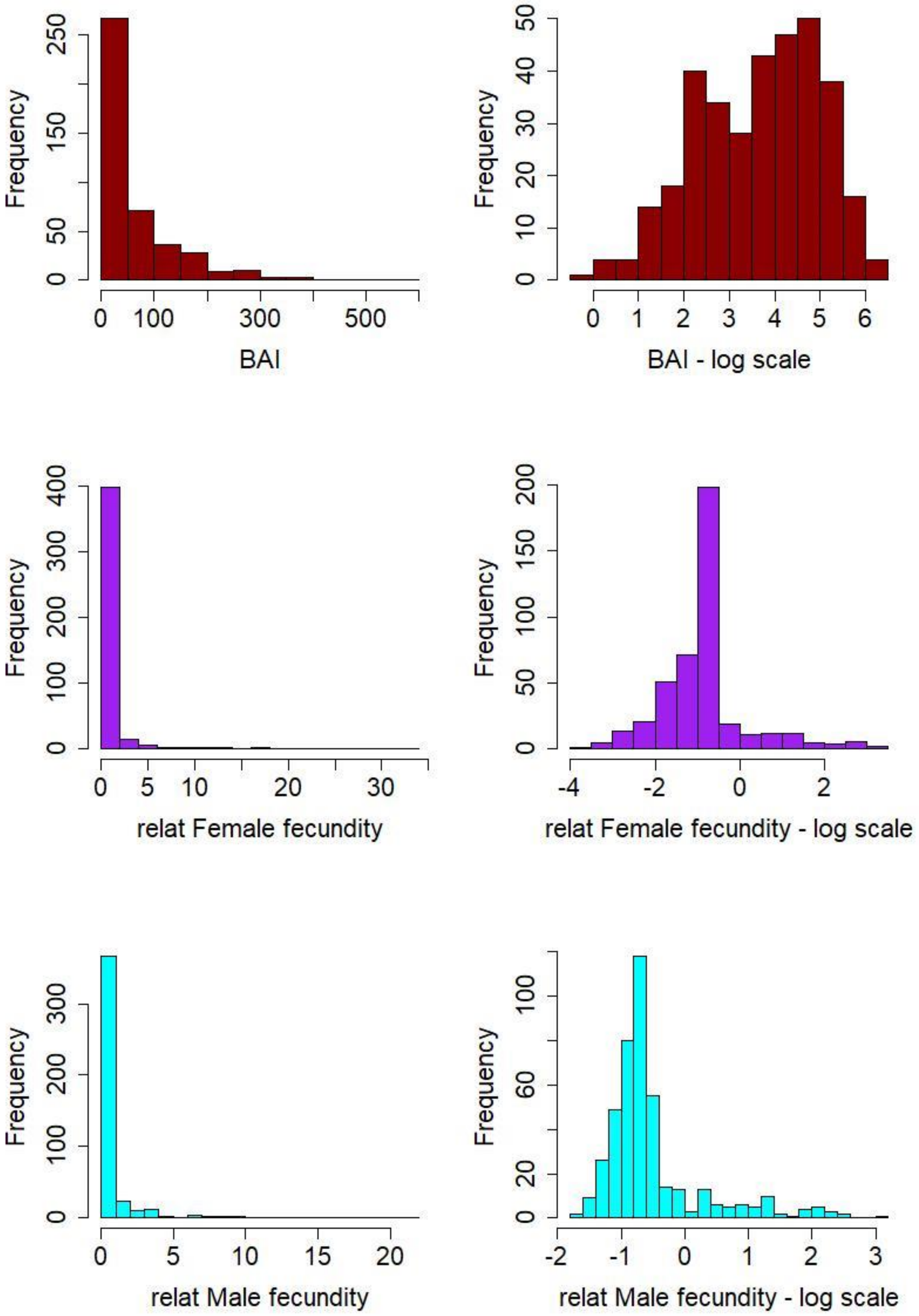
Frequency
BAI
BAI - log scale
relat Female fecundity
relat Female fecundity - log scale
relat Male fecundity
relat Male fecundity - log scale

C- Basal area Increment (BAI), equation 3

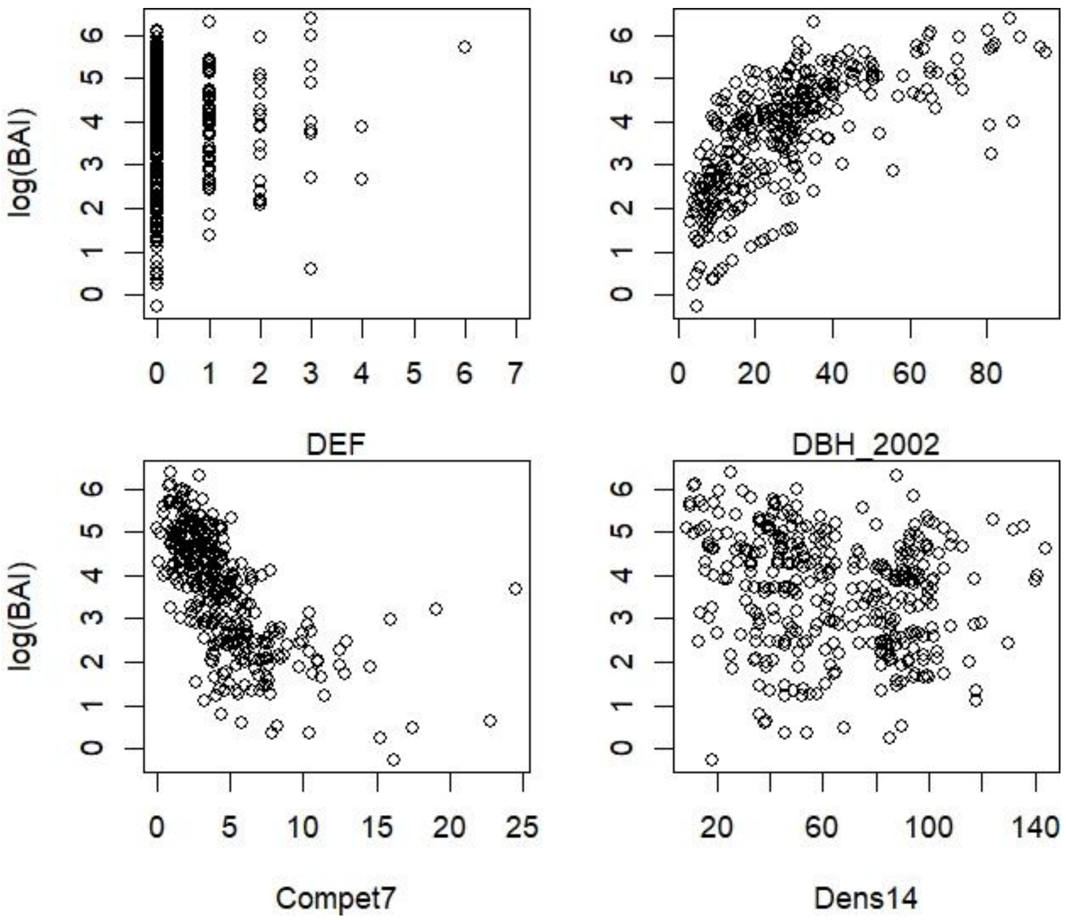

C- Female fecundity, equation 3

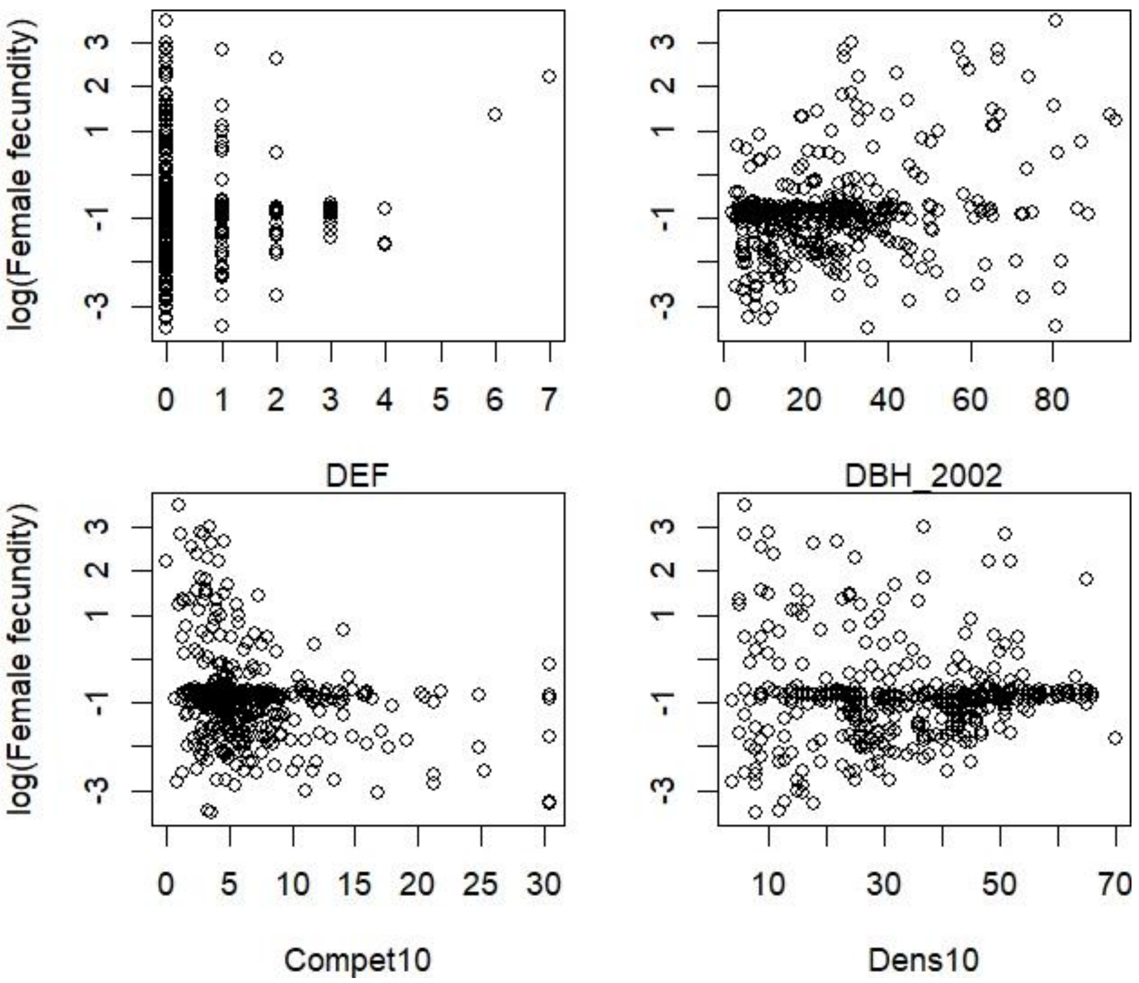

Note that the denser line of point on the female fecundity scatter plots correspond to individual female fecundity values estimated around the mean female fecundity. In other terms, these are individuals for which the dataset does not contain enough information to estimate fecundity. They should however not bias the linear model, even though they likely decrease the effective number of degree of freedom.

The scatter plots at the bottom explain why Comp10 has a negative effect whereas Dens10 has a positive effect on female fecundity. These opposite Type III effects of competition and density are probably driven by the facts that (1) only trees with low competition indexes showed a high female fecundity and that (2) only trees with low density in the neighborhood showed a very weak female fecundity. Moreover, the positive correlation between compet10 and Dens10 may also contribute to these effects (cor=0.10, pval=.02).

C- Female fecundity, equation 4

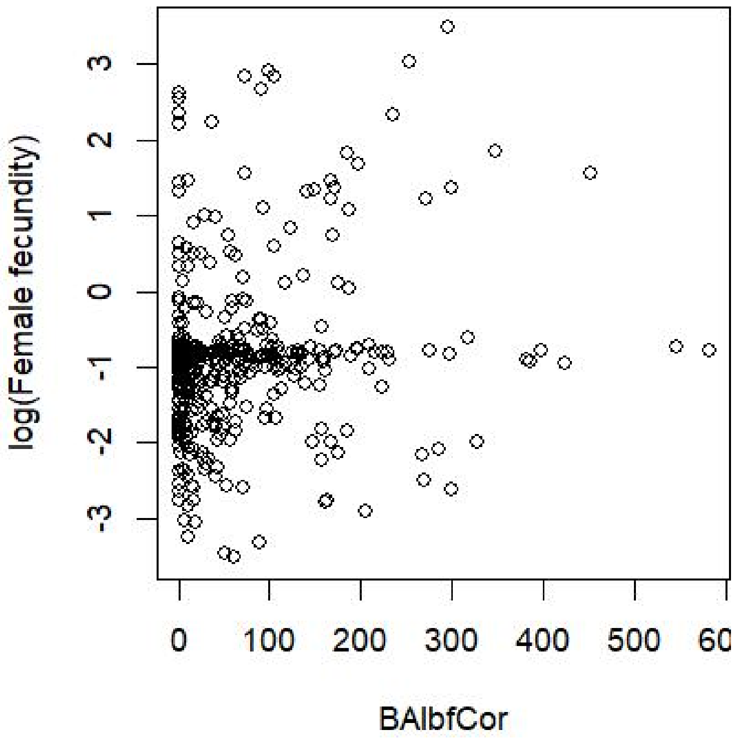

C- Male fecundity, equation 3

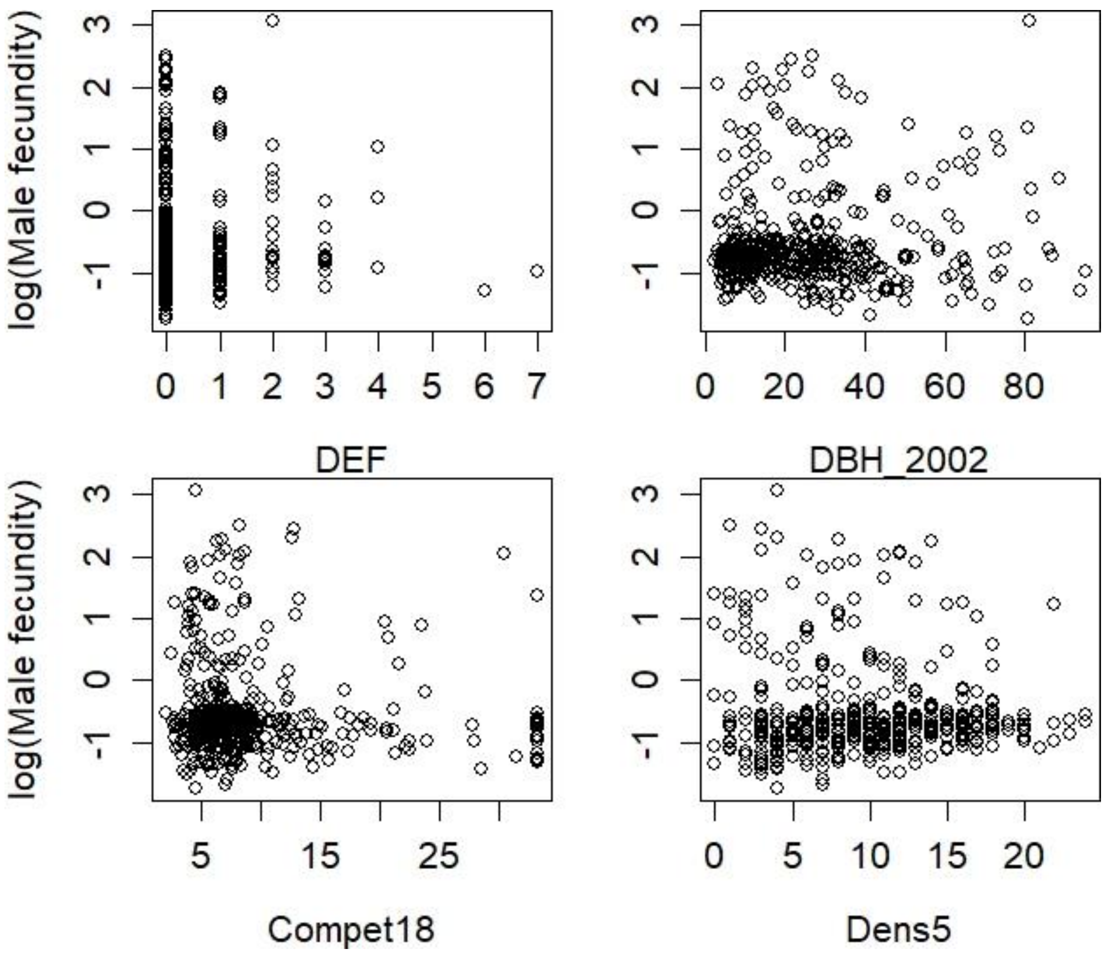

### Figure S5: Effect of size and competition on defoliation.

We used a model similar to equation (3) to investigate the effects of tree size and competition on defoliation : $DEF = DBH_{2002} + Compet_{dmax} + Dens_{dmax} + DBH_{2002}{:}Compet_{dmax} + DBH_{2002}{:}\ Dens_{dmax}$

**A. Analysis of variance table of the model:** the adjusted $R^2$ was 0.28. For each term, we give the type III sum of squares (SSQ) and degree of freedom (df), and for each predictor, the estimate of its effect, the standard error (S.E.) and associated t and p-value. Variance inflation factors (VIF) were computed with R package CAR.

| | SSQ | df | Estimate | S.E. | t | P-value | VIF |
|---|---|---|---|---|---|---|---|
| **$DBH_{2002}$** | 5.24 | 2 | -6.541 | 2.143 | -3.052 | 0.002 | 2.66 |
| **$DBH^2_{2002}$** | | | 0.506 | 1.805 | 0.280 | 0.780 | |
| **Compet19** | 1.77 | 1 | 0.011 | 0.006 | 1.891 | 0.059 | 1.22 |
| **Dens20** | 10.86 | 1 | 0.003 | 0.001 | 4.682 | 0.000 | 1.26 |
| **$DBH_{2002}$:Compet19** | 16.06 | 2 | 0.516 | 0.105 | 4.893 | 0.000 | 1.78 |
| **$DBH^2_{2002}$: Compet19** | | | 0.176 | 0.089 | 1.973 | 0.049 | |
| **$DBH_{2002}$:Dens20** | 16.44 | 2 | 0.109 | 0.020 | 5.407 | 0.000 | 2.55 |
| **$DBH^2_{2002}$: Dens20** | | | 0.016 | 0.016 | 1.035 | 0.301 | |
| **residuals** | 209.50 | 423 | | | | | |

**B. Interaction plot** showing regression lines of defoliation against DBH for three levels of B1-competition or B2-density, corresponding to +/- 1 standard deviation from the mean. Confidence interval at 80% are displayed around each regression line.

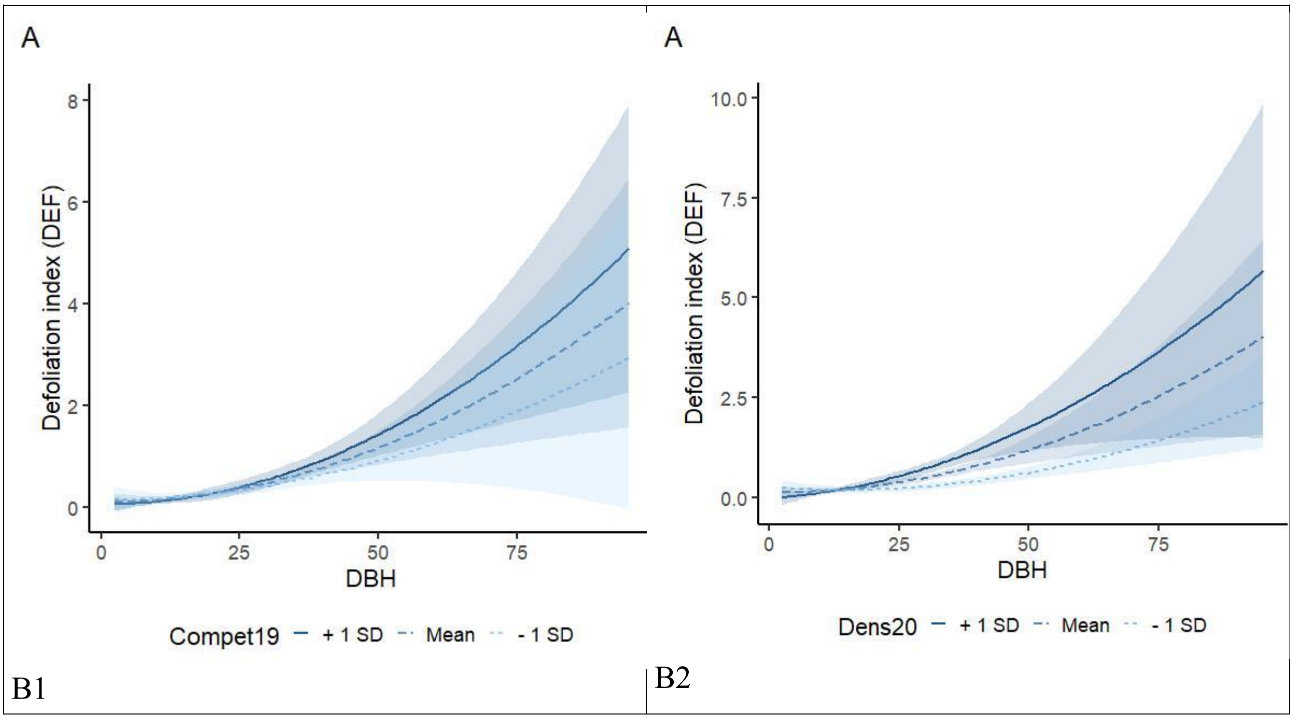

**Figure S6: Relationship between growth estimated from ring-width and growth estimated from inventory data.**

The graph on the left plots the cumulated radial growth from 2002 to 2012 respectively estimated from ring-width (x-axis) and inventory (y-axis). The graph on the right plots the cumulated basal area increment from 2002 to 2012 (BAI) respectively estimated from ring-width (x-axis) and inventory (y-axis), based on the 90 cored trees. The correlation between estimates in showed on each graph.

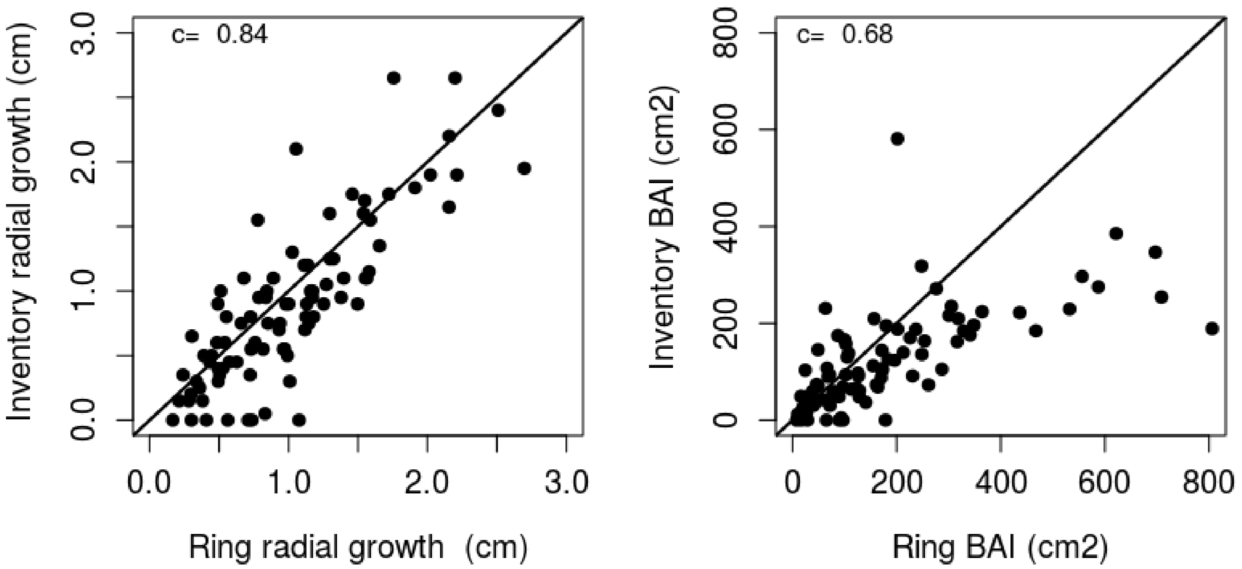

**Figure S7: Diagnostic plot for the linear regression model described by equation 3 and the three response variables: A: log(BAI; B. log(F♀) and C:log(F♂)**

A.

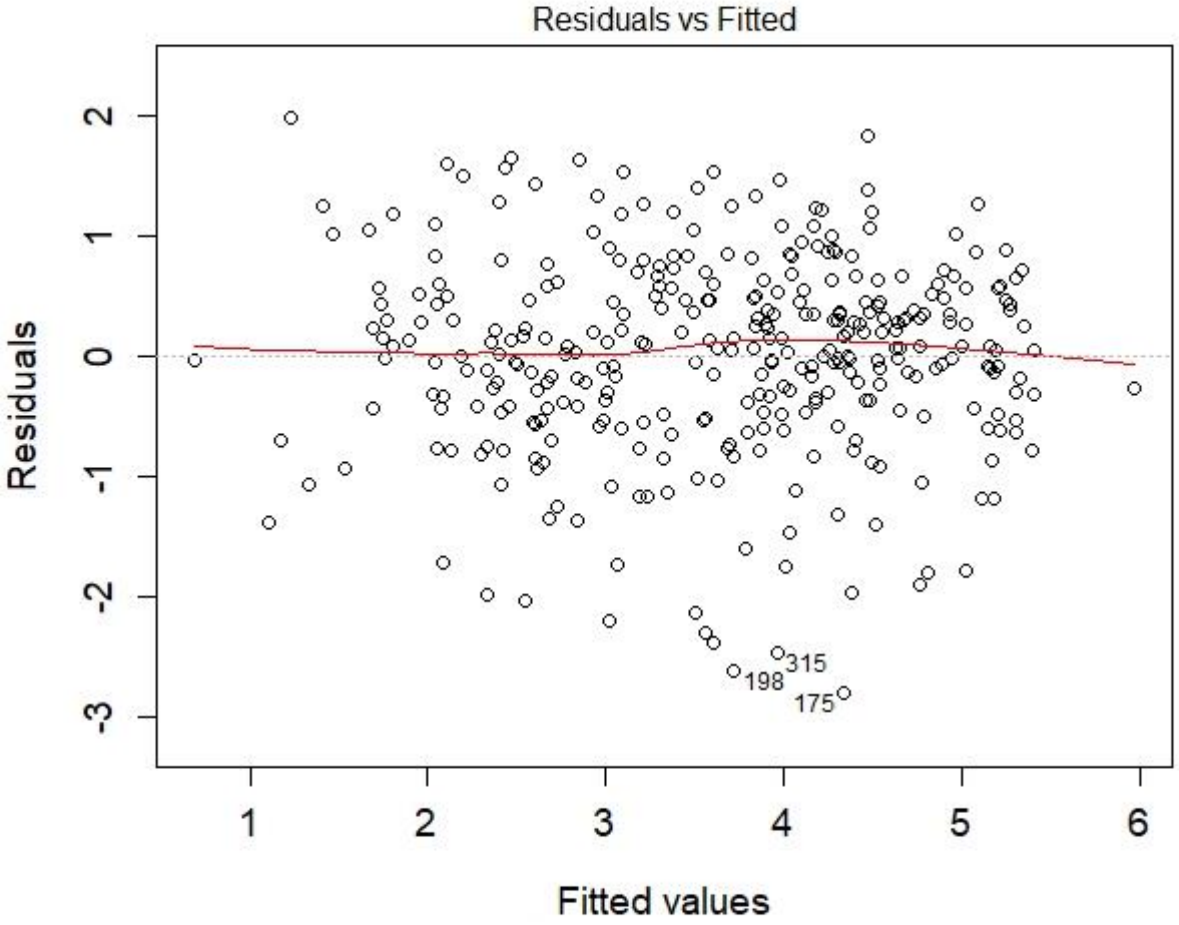

B

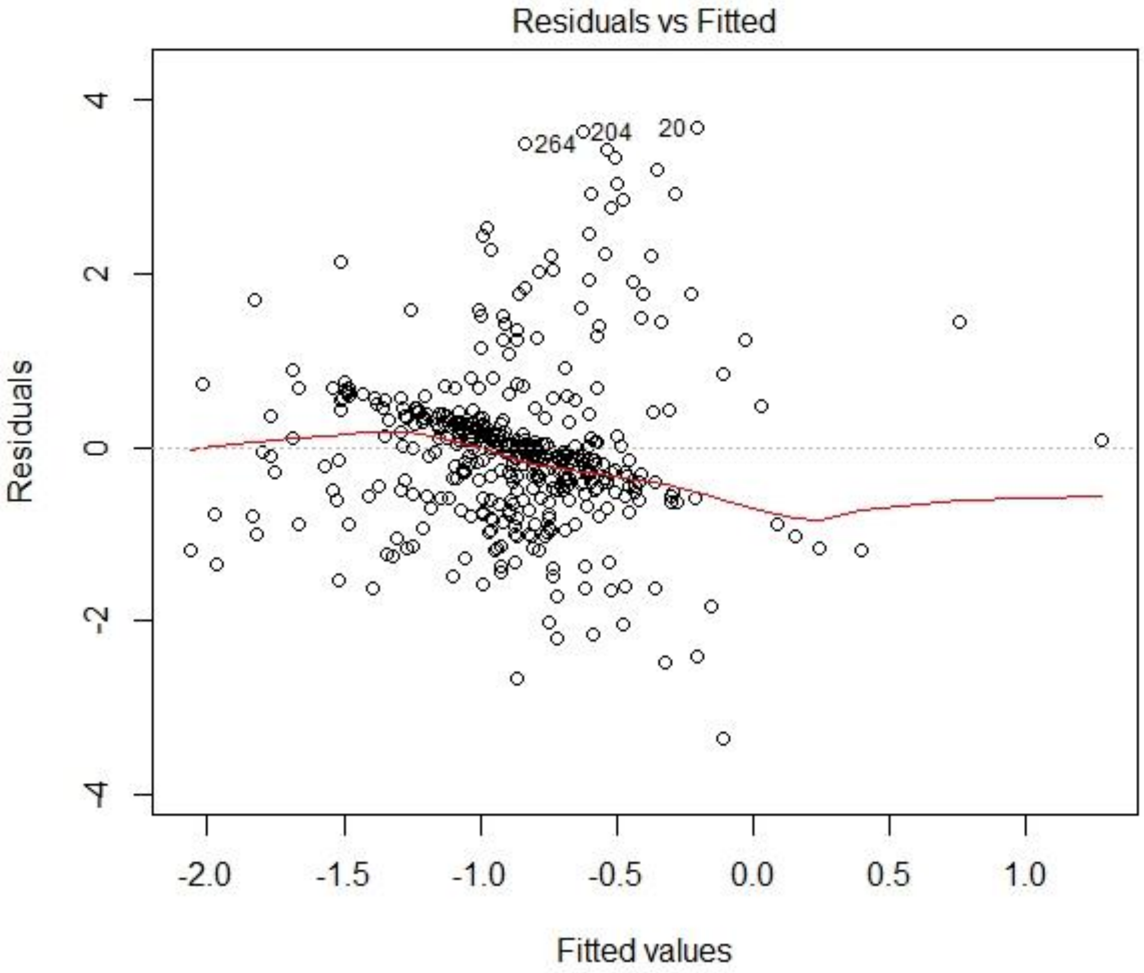
Residuals vs Fitted
264
204
20
4
2
0
-2
-4
Residuals
-2.0
-1.5
-1.0
-0.5
0.0
0.5
1.0
Fitted values

C

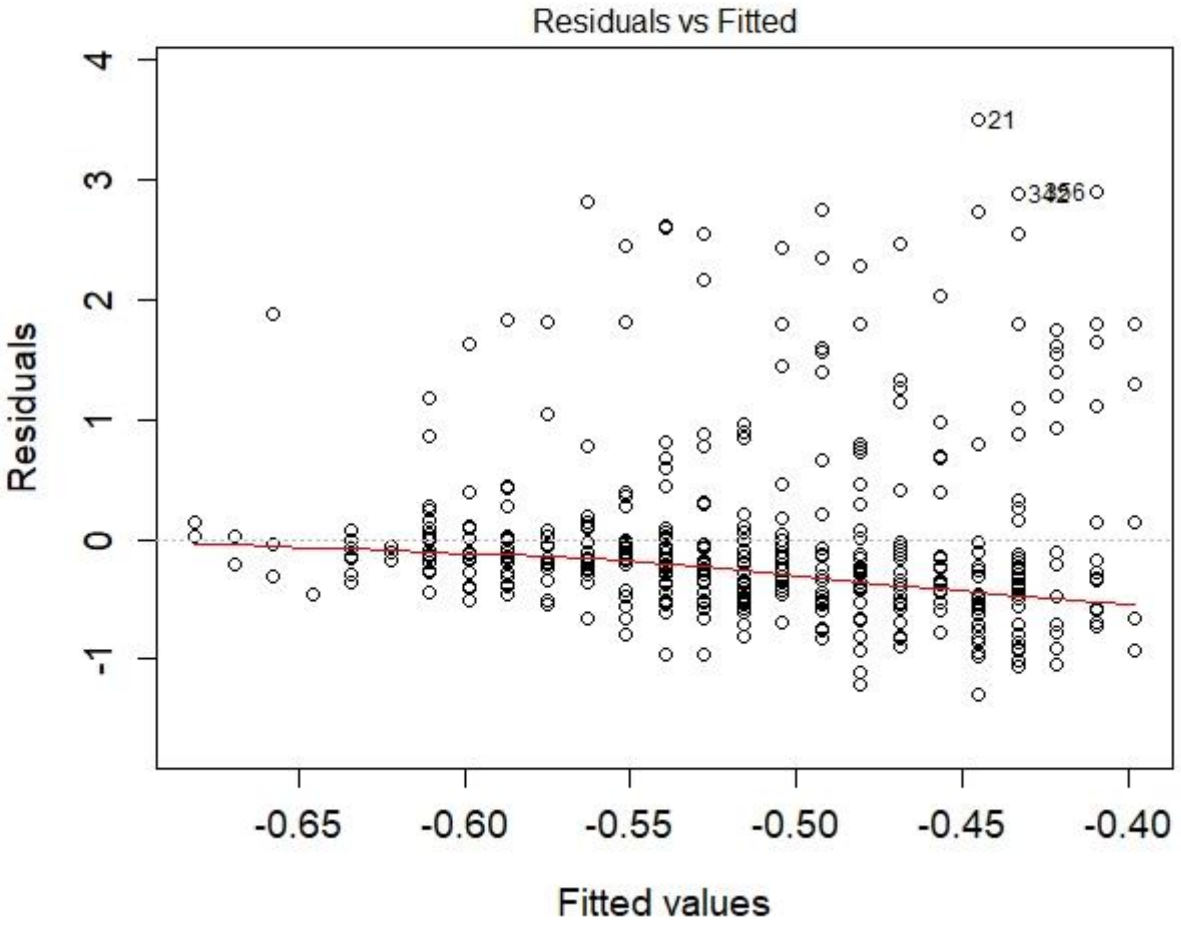
Residuals vs Fitted
21
Residuals
4
3
2
1
0
-1
-0.65
-0.60
-0.55
-0.50
-0.45
-0.40
Fitted values

**Figure S8: Interaction plots for DEF , BAI, and $DBH_{2002}$ effects on female fecundity.** Regression lines are plotted for three values of each moderator variable, corresponding to +/- 1 standard deviation from the mean. Confidence interval at 80% are shown around each regression line. Points are the observations.

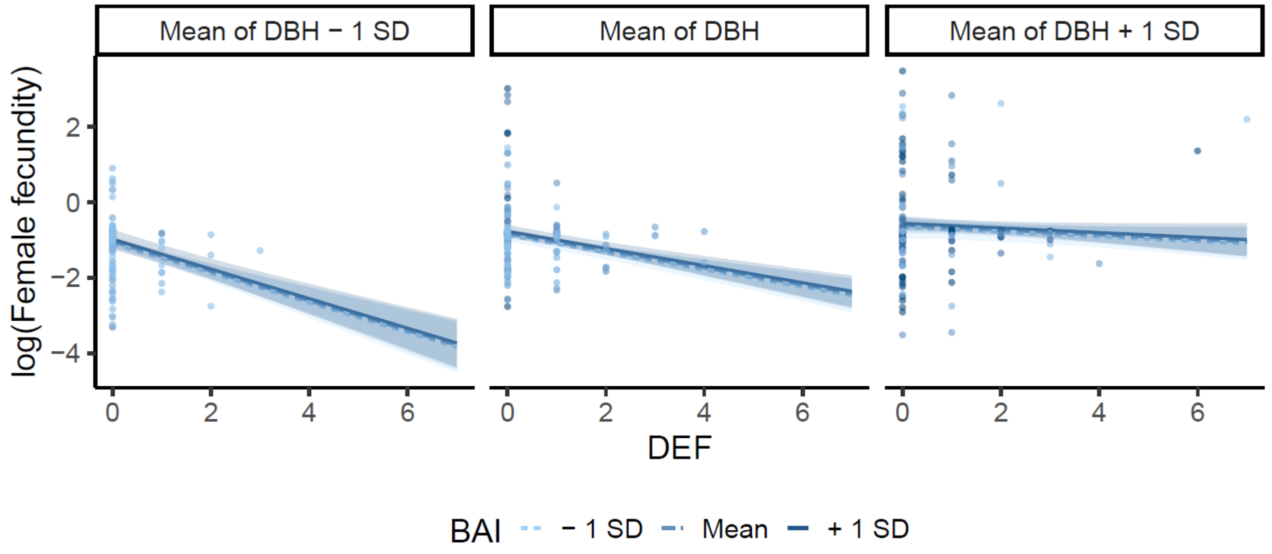

**Figure S9: Diagnostic plot for the linear regression model described by equation 4**

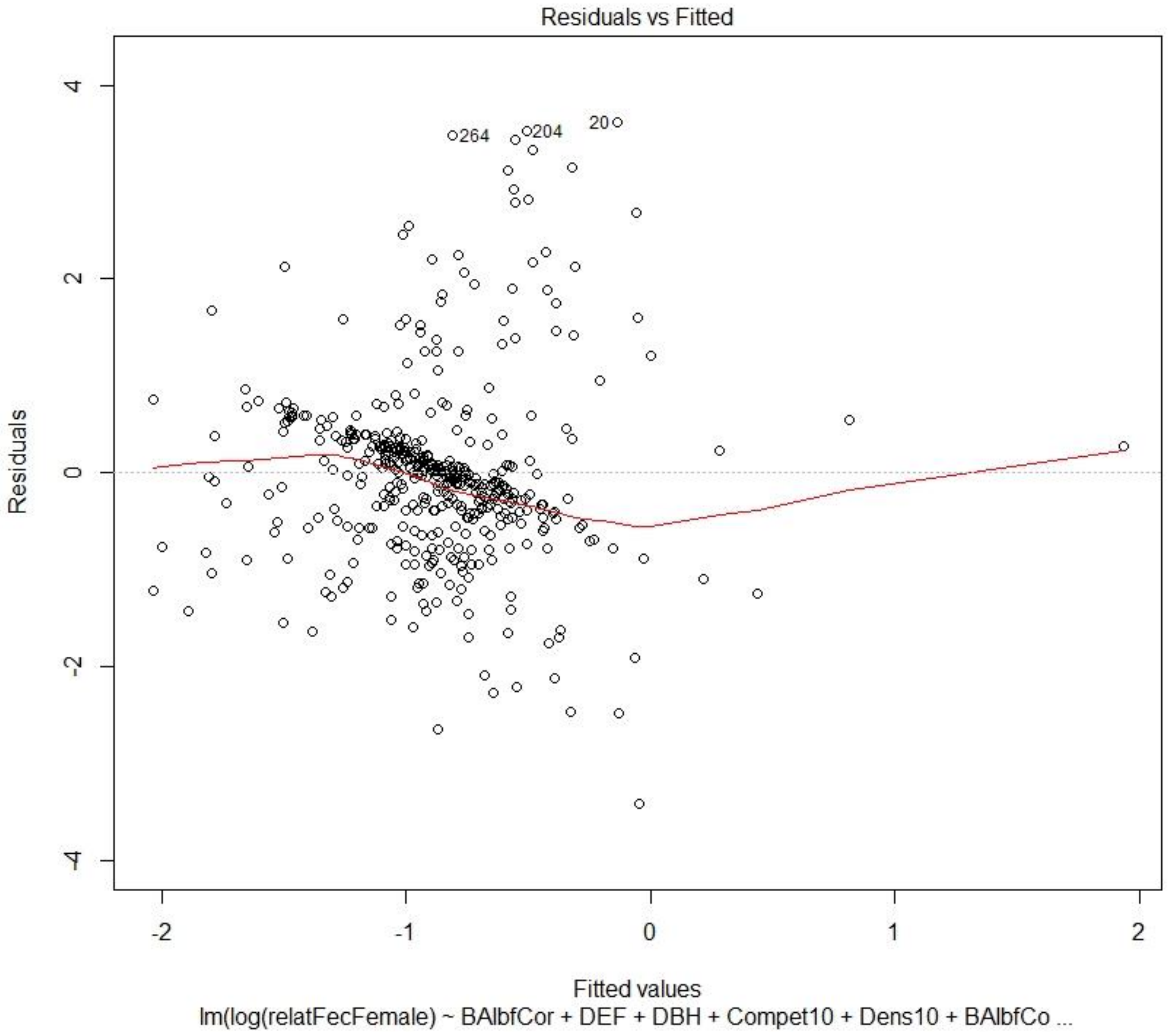
